# Germinal center-independent memory B cells provide rapid protection from lethal influenza challenges

**DOI:** 10.64898/2026.07.15.738751

**Authors:** Elif Çakan, Faez Amokrane Nait Mohamed, Anson Hui, Essi C. Logan, Nicole Ingram, Caroline A. Alexander, Jack Smerczynski, Chinmay Varma Gadiraju, Emma Barnes, Daniel A. Mounier, Bruce D. Walker, Daniel Lingwood, Shiv Pillai

## Abstract

Memory B cell recall responses are crucial for rapid protection from pathogens expressing previously encountered antigens. While germinal centers (GCs) contribute to durable B cell memory in many contexts, GC formation is attenuated or completely abrogated during some severe infections. Whether GC-independent responses generate functional memory B cells that may contribute to protective immunity remains unclear. Using mice lacking GCs, we identified a durable class-switched GC-independent memory B cell population that was generated dominantly through a T-cell-dependent response. Vaccine-induced non-GC memory B cells demonstrated greater diversity and provided humoral protection from matched and diverse vaccine-unmatched influenza viral challenges. These results identify a unique, durable, diverse, GC-independent memory B cell population that can mediate rapid protection from severe infections by mutable pathogens.

## Introduction

Germinal centers (GC) are the sites where durable, high-affinity memory B cells are generated. When B cells that have received help from T cells enter GCs, they undergo iterative cycles of somatic hypermutation (SHM) of antibody genes and selection to generate durable, high-affinity memory B cells (*1–3*). The initial T cell-B cell interactions that occur at the T-B border may induce class switching to many isotypes well before the formation of GCs (*4*, *5*).

Many severe intracellular infections in animal models and humans result in the loss of GCs (*C*–*8*), but in these contexts, class-switched B cell responses are preserved (*S*). Diverse IgM or class-switched, T-independent and T-dependent responses that are induced before GC formation, or in the absence of GCs, are widely referred to as “extrafollicular” B cell responses (*10–13*). While extrafollicular B cell populations have been widely studied in inflammatory or autoimmune contexts, few studies have explored whether robust and protective class-switched memory B cells can be generated in the absence of GCs (*12*, *14–17*). Mice without GCs have been shown to generate neutralizing IgG antibodies after immunization with the SARS-CoV-2 Spike antigen (*18*). Antibody-dependent protection against a homologous viral challenge after vaccination with inactivated influenza virus has also been observed in mice similarly engineered to lack GCs (*1S*). However, the identity of the memory B cell subsets responsible for this form of humoral immunity remains to be established, and whether these B cells are durable and capable of providing rapid protection from future infections is also not known.

Here, we utilized an established mouse model genetically engineered to lack T follicular helper (Tfh) cells, and therefore lacking GCs, to define the features and functions of class-switched B cells, including distinct durable memory B cell populations that are either GC-derived or that can emerge independently of Tfh cells and GCs (*20*). The non-GC memory B cell population exhibits greater diversity and therefore greater potential immunological breadth than GC-derived memory B cells. We show that vaccine-induced non-GC B cell responses are able to protect against lethal doses of homologous and heterologous influenza viruses. The abundance of influenza-specific non-GC memory B cells correlated with survival after a lethal challenge in both control mice and mice lacking GCs, and adoptive transfer of these memory B cells conferred protection against death in the absence of any other B cells. These results suggest that vaccine strategies that facilitate the induction of diverse, class-switched GC-independent memory B cells may facilitate the generation of rapid protective responses against severe viral pathogens that have a propensity to generate variants that can evade previously generated neutralizing antibodies.

## Results

### Class-switched non-GC memory B cells have a distinct trancriptome and can be identified by specific cell surface markers

To study class-switched non-GC B cell responses, we analyzed Bcl6^fl/fl^CD4^Cre^ mice, which lack T follicular helper (Tfh) cells, for which Bcl6 is a defining transcription factor (*21*). Tfh cells are essential for GC formation, and the absence of Tfh cells leads to the lack of GCs in this model.

We immunized control (Bcl6^fl/fl^) and Bcl6^fl/fl^CD4^Cre^ mice with a previously described protein antigen (HIV gp140(*22*)) and showed that class-switched and antigen-specific splenic B cells were only partially depeleted in the absence of germinal centers (Fig. 1A). We used the GL7 antibody, which recognizes a sialic acid species found on activated B cells, to broadly distinguish activated B cells from resting putative memory B cell populations (*23*). Of note and as discussed later in the manuscript, we refer to the GL7^-^ class-switched resting GC-populations and non-GC populations as memory B cells since these populations persist at day 90 and day 180 after immunization. We categorized GL7^+^ B cells as activated B cells (GL7^+^CD73^-/low^), or as GC B cells (GL7^+^CD73^+^).

**Figure 1.**
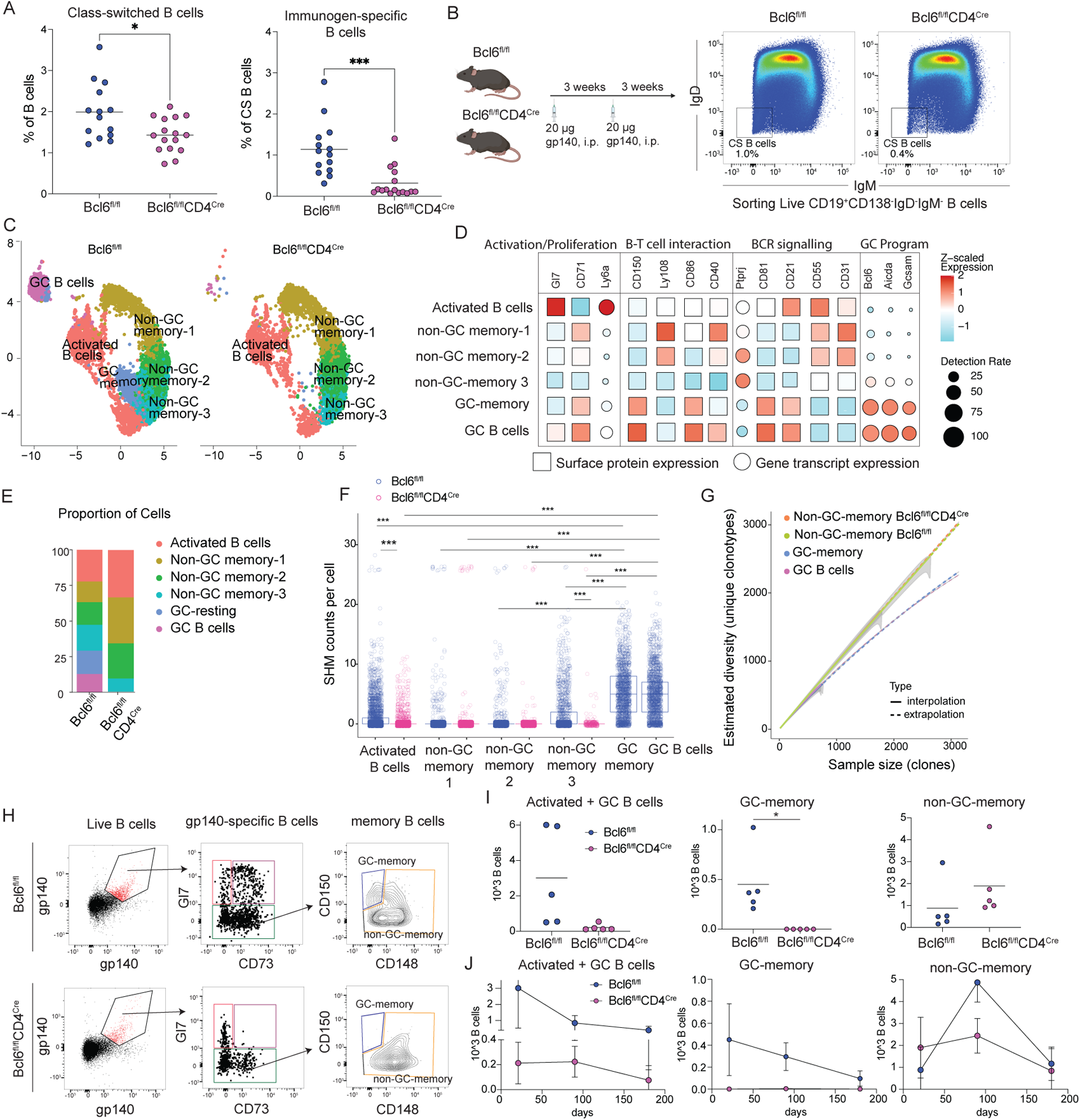
GC-independent class-switched B cells with low SHM frequency and high BCR diversity, contribute to durable B cell responses. (A) Frequencies of IgD^-^IgM^-^ (class switched) and gp140-specific B cells in Bcl6^fl/fl^CD4^Cre^ (n=16) and Bcl6^fl/fl^ (n=14) mice after 2 immunizations with gp140. Data are aggregated from 3 independent experiments. (B) Experimental plan for IgD^-^IgM^-^ B cell sorting from Bcl6^fl/fl^ (n=5) and Bcl6^fl/fl^CD4^Cre^ (n=5) mice 3 weeks after the gp140 boost. (C) UMAP (uniform manifold approximation and projection) based clustering of IgD^-^IgM^-^ B cells based on surface expression using CITE-seq (Cellular Indexing of Transcriptomes and Epitopes by sequencing). (D) Dot plots showing the average expression and the percentage of the cells expressing the indicated genes or surface markers in Bcl6^fl/fl^ mice; (E) Frequencies of indicated clusters in Bcl6^fl/fl^ (n=5) and Bcl6^fl/fl^CD4^Cre^ (n=5) mice. (F) SHM counts per cell of each B cell cluster in Bcl6^fl/fl^ and Bcl6^fl/fl^CD4^Cre^ mice. (G) Rarefaction curves showing the diversity of BCRs for T cell dependent non-GC and GC B cells. Each curve represents the indicated cluster in Bcl6^fl/fl^ (n=5) or Bcl6^fl/fl^CD4^Cre^ (n=5) mice. (H) Gating strategy for dissecting gp140-specific B cells into activated, GC, GC-memory and non-GC memory clusters using the cluster-defining markers generated from the surface expression library. (I) gp140-specific B cell numbers for each indicated B cell subset at 3 weeks (n=5), (J) Changes in gp140-specific B cell numbers at 3 weeks (n=5), 3 months (n=5) and 6 months (n=5) after the second dose of gp140 immunization in Bcl6^fl/fl^ and Bcl6^fl/fl^CD4^Cre^ mice. The dots represent the mean of cell numbers for each population with standard deviation. Data are aggregated from two independent experiments.

To further characterize class-switched (IgD^-^IgM^-^) B cells, we sorted these cells from the spleens of Bcl6^fl/fl^ control and Bcl6^fl/fl^CD4^Cre^ mice for scRNA-seq, CITE-seq and BCR-seq library preparation (Fig. 1B). Using CITE-seq libraries, B cells were grouped into 6 clusters that included 4 GC-independent and 2 GC-dependent populations (Fig. 1C, E). The UMAP based clustering of B cells was very similar in each of the control mice (n=5), as well as in the Bcl6^fl/fl^CD4^Cre^ mice (n=5) (Supp. Fig. 1A). The activated B cell cluster expressed high levels of *ApoE* and *FCRL5*, genes known to be induced in activated B cells (Supp. Fig. 2A). In addition to the lack of GC B cells, a GL7^-^ memory B cell cluster was missing in the absence of GCs (Fig. 1C). This GC-dependent class-switched memory B cell cluster was characterized as CD150^high^CD148^-^GL7^-^ B cells, expressing high levels of genes associated with the GC program including *Aicda*, *BclC* and *Gcsam* as well as the surface expression of CD150 *(SLAM)* (Fig. 1D, Supp. Fig. 1B-D, Supp. Fig. 2). In contrast, class-switched non-GC memory B cells expressed high levels of CD148 (encoded by *Ptprj),* a receptor-type tyrosine phosphatase, previously reported to be involved in BCR activation (*24*) (Fig. 1D, Supp. Fig.1B-D).

Three of the clusters represented non-GC memory B cells. These exhibited lower SHM frequencies compared to GC-derived B cell populations, further supporting their non-GC origin (Fig. 1F, Supp. Fig. 3A-B). In addition, non-GC-memory B cells had more diverse BCRs compared to GC-memory B cells, consistent with lower levels of GC-related clonal expansion and affinity maturation (Fig. 1G). BCR connectivity analyses revealed that the “activated B cell” cluster is a heterogeneous population connected to both GC-dependent and independent B cells, explaining their higher SHM frequencies in Bcl6^fl/fl^ control mice and consistent with a previous report (*25*) (Supp. Fig. 3D, Fig. 1F). In contrast, GC B cells and GC memory B cells shared the majority of their common BCRs with each other. The three non-GC memory B cell clusters were connected mostly to activated B cells and to one another (Supp. Fig. 3D). Non-GC B cells had significantly lower kappa/lambda light chain ratios and differences in V(D)J usage compared to GC B cells and GC-memory B cells (Supp. Fig. 3C-I).

The data show that class-switched GC-memory and non-GC memory B cells are associated with distinct transcriptional programs and surface marker phenotypes, and can be reliably differentiated based on cell surface expression of CD150 and CD148 in resting (GL7^-^) class-switched B cells.

#### Class-switched non-GC-memory B cells are durable

Using CD150 and CD148 as markers based on their differential surface expression in class-switched GC-dependent (CD150^high^CD148^-^) and GC-independent (CD150^-^CD148^+^) memory B cells, we established a flow cytometry gating strategy to identify memory B cell clusters of GC-independent origin (Fig. 1H and Supp. Fig 4A). Using this strategy, we showed that class-switched GC-dependent memory B cells were depleted in the absence of GCs, validating our markers (Fig. 1I). Furthermore, three class-switched non-GC memory B cell clusters could be distinguished by their levels of CD24, CD73, CD31 and Slamf6 expression, which allowed us to further dissect these non-GC memory B cell subsets (Supp. Fig. 4A).

Next, we phenotyped gp140-specific B cells in the spleen 3 weeks, 3 months, and 6 months after immunization to investigate whether antigen-specific class-switched GC-memory and non-GC-memory B cells differ in terms of durability. GL7^+^ B cell populations were abundant at day 21 and then rapidly declined, but GC-memory B cells declined far more slowly than GC B cells, supporting that they are memory B cells (Fig. 1J, Supp. Fig. 4B). At later time points, the vast majority of antigen-specific class-switched B cells were non-GC-memory B cells. Both gp140-specific GC-memory and non-GC-memory B cells persisted at 6 months after immunization (Fig. 1J, Supp. Fig. 4B). These results indicate that both GC and non-GC-derived class-switched memory B cells may contribute to durable B cell responses.

Absolute numbers of class-switched non-GC memory B cells peaked at the 90-day post-immunization time point, and these numbers were distinctly higher in control littermates compared to Bcl6^fl/fl^CD4^Cre^ mice (Fig. 1I, J). These data indicate that although non-GC class-switched memory B cells do not *require* Tfh cells or GCs for their development or maintenance, T-B collaboration that generates these memory B cells not only involves non-Tfh helper T cells but can also involve Bcl-6-expressing Tfh cells that have not yet entered the GC, consistent with previous reports regarding Tfh cell participation in T-B collaboration at the T-B border (*2C*–*28*).

#### Class-switched non-GC memory B cells contribute to recall responses

To confirm that similar class-switched memory B cells emerge in response to a range of protein antigens and to determine how class-switched GC and non-GC memory B cells respond upon reactivation by the same antigen, we immunized wild-type mice with 2 doses of the model T-dependent antigen nitrophenyl-keyhole limpet hemocyanin (NP-KLH) and separately sorted total class-switched GC memory B cells, non-GC memory B cells and CXCR5^hi^PD1^hi^ Tfh cells. Individual memory B cell populations and Tfh cells were transferred into Rag2 KO mice that lack mature B and T cells. One day after cell transfer, the recipient mice were immunized with NP-KLH (Fig. 2A).

**Figure 2.**
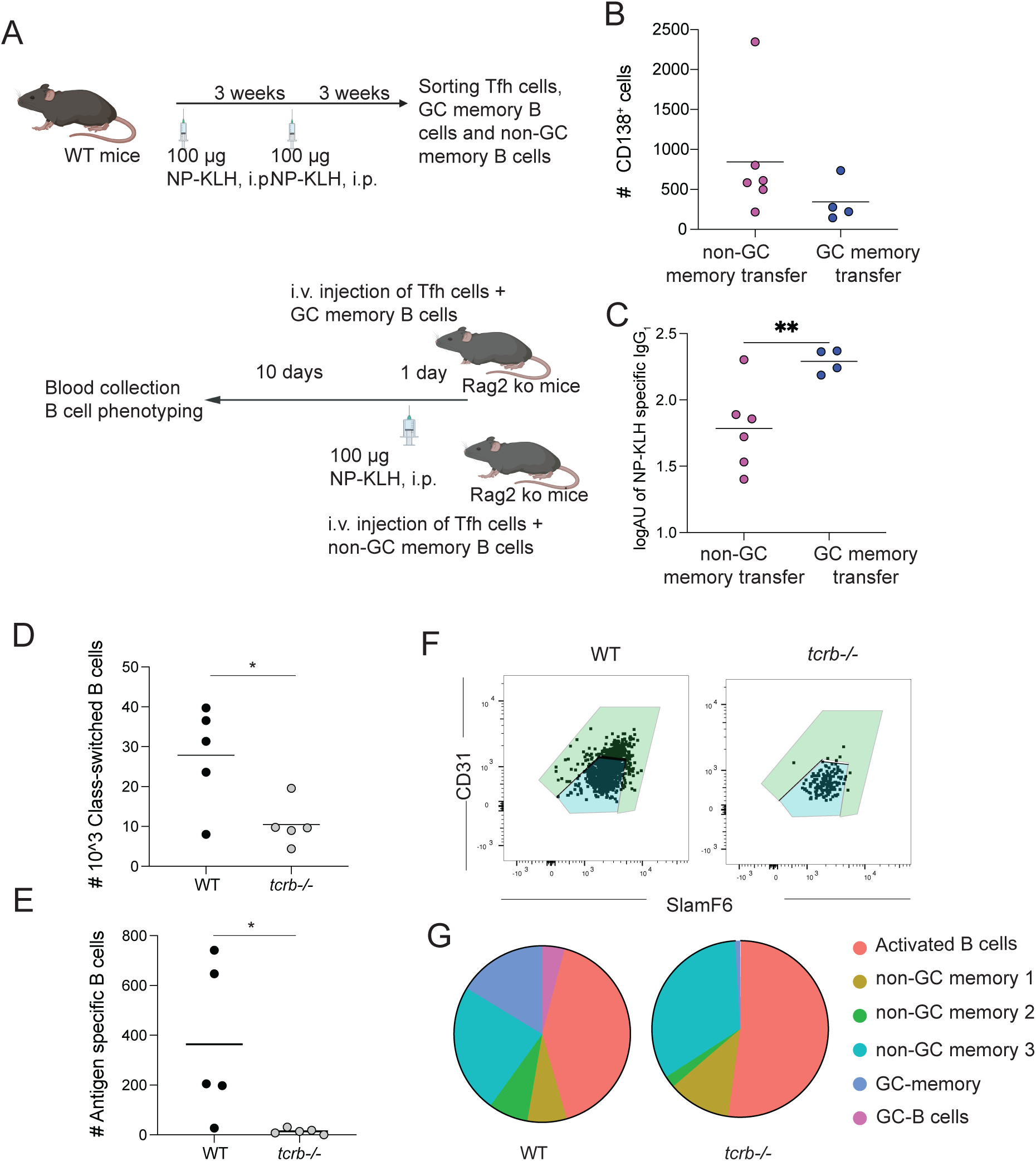
Class-switched non-GC memory B cells contribute to memory recall responses that are primarily linked to T-cell dependent immune reactions. (A) GC-memory and non-GC memory B cells were sorted from WT mice (n=60) after two doses of NP-KLH and separately transferred into Rag2ko mice (n=6 for non-GC memory B cell transfer, n=4 for GC-memory B cell transfer) along with sorted Tfh cells. Rag2ko mice were immunized with NP-KLH the day after cell transfer. (B) Absolute numbers of CD138^+^ B cells in the spleen, (C) NP-KLH specific IgG_1_ levels in the serum 10 days after NP-KLH immunization. Absolute numbers of class switched B cells (D) and class-switched antigen-specific B cells (E) after 2 doses of 20 µg gp-140 immunization in WT (n=5) and *tcrb-/-* (n=5) mice. (F) Gating strategy to distinguish T cell dependent and independent non-GC memory B cells using antibodies to CD31 and SlamF6. (G) Frequencies of each B cell cluster in WT and *tcrb-/-* mice shown as pie charts. Frequencies were calculated as the average of the frequency for each cluster in 5 mice for each group.

Phenotyping B cells from the spleen 10 days after immunization revealed that both GC and non-GC memory B cell populations were reactivated and differentiated into CD138^+^ plasmablasts (Fig. 2B). Consistent with the differentiation of memory B cells into antibody secreting cells, immunization of both transferred memory B cell populations resulted in NP-KLH-specific IgG secretion, most prominently to antibodies of the IgG_1_ subtype (Fig. 2C, Supp. Fig. 6A). We conclude that both GC and non-GC memory B cells upon reactivation can give rise to plamablasts and thereby contribute to recall responses.

#### The majority of class-switched non-GC memory B cells are T cell-dependent

To explore whether non-GC memory B cell populations are linked to T cell help, we immunized T cell receptor (TCR) beta knock out (*tcrb*-/-) (n=5) and wild-type (n=5) mice with 2 doses of gp140 and phenotyped splenic B cells. As expected, total class-switched and antigen-specific class-switched B cells were markedly reduced in the spleens of *tcrb*-/- mice compared to wild-type mice (Fig. 2D-E). The numbers of all GC-dependent and GC-independent memory B cell subsets were depleted in *tcrb*-/- mice. Interestingly, whereas most clusters of non-GC memory B cells were largely or significantly dependent on T cell help, one specific sub-population of non-GC memory B cells, (CD31^hi^Slamf6^hi^ non-GC memory B cells) could not be identified in the absence of T cell help, indicating an absolute requirement for T cell help in this class-switched non-GC memory B cell subset (Fig. 2F-G).

These results indicate that the class-switched non-GC B cell responses are heavily T cell dependent. Although it has been argued that Bcl6-expressing Tfh cells are necessary for extrafollicular reactions, many other studies, including ours, demonstrate that non-Tfh CD4+ T cells can induce class-switched B cell responses in the absence of Tfh cells. While Tfh cells are not required for non-GC memory B cell generation, they can contribute in part to the accumulation of cells in the non-GC memory B cell compartment (*2C*, *2S*).

#### Vaccine-induced non-GC humoral immunity provides protection from both homologous and heterosubtypic inffuenza challenges

To evaluate the potential protective activity of the non-GC memory B cell compartment, we first performed 2x sequential immunization with recombinant influenza hemagglutinin (HA) H3 trimer ectodomain [A/Hong Kong/1/1968 (HK68)] (plus Sigma adjuvant) followed by lethal challenge with strain-matched H3N2 X-31 virus within Bcl6^fl/fl^CD4^Cre^ and Bcl6^fl/fl^ littermate mice (Fig. 3A,B). The vaccine regimen was protective in the presence and absence of GCs [Bcl6^fl/fl^CD4^Cre^ mice (6 of 9) and Bcl6^fl/fl^ mice (10 of 10), each relative to non-immunized control (0 of 10), P<0.0077 and P<0.0001, respectively (Mantel-Cox test of survivorship)] (Fig. 3C-D).

**Figure 3.**
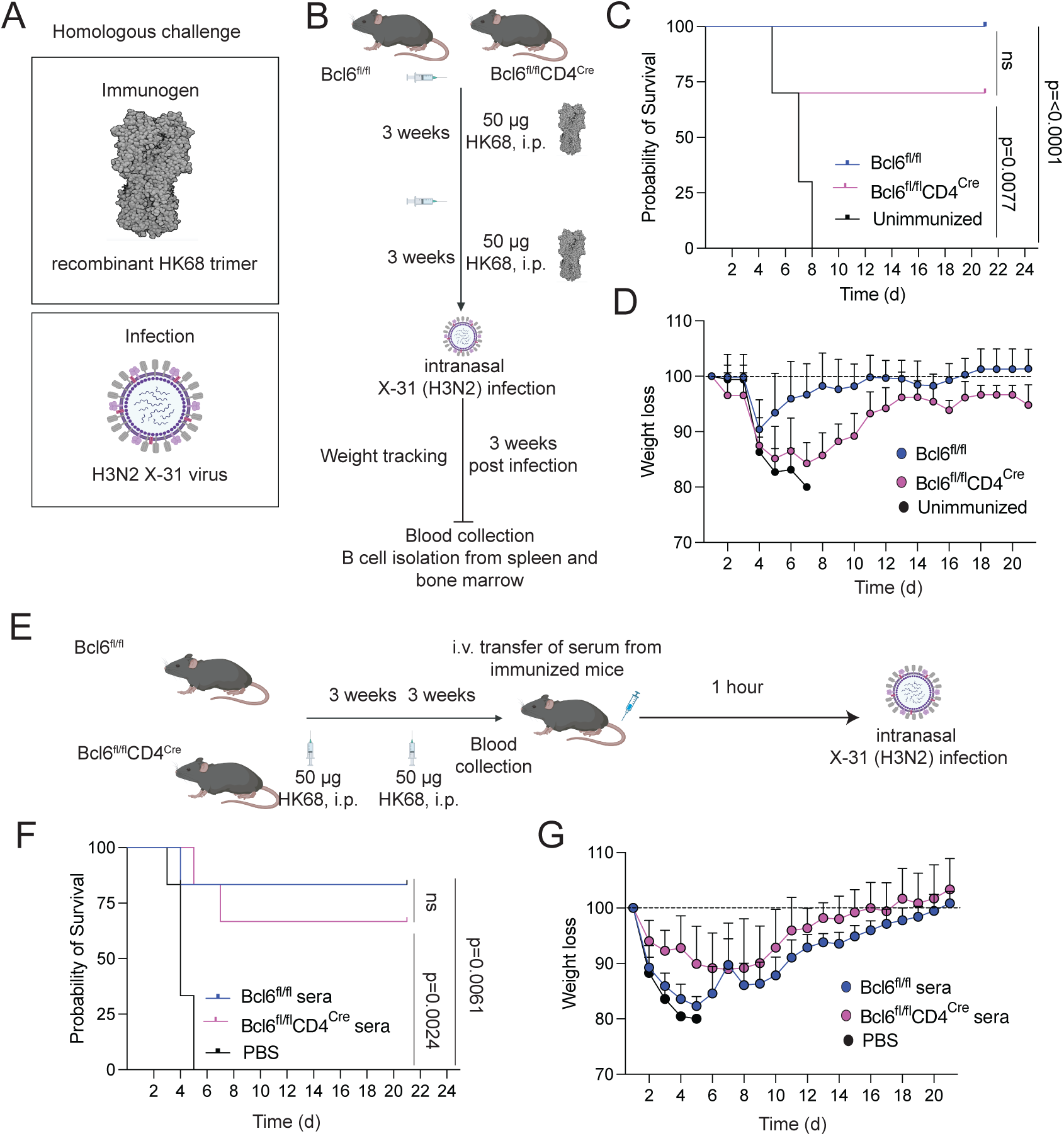
Non-GC derived humoral immunity protects against a lethal homologous influenza challenge. (A, B) 3 weeks after 2^nd^ dose of 50 µg HK68 immunization, Bcl6^fl/fl^ (n=10) and Bcl6^fl/fl^CD4^Cre^ (n=10) mice were lethally challenged with X-31 (H3N2) virus and monitored up to 21 days post infection. (C) Kaplan-Meier graph showing the probability of survival and (D) percentage of weight loss after lethal X-31 challenge. (E) Experimental plan for the passive transfer of sera from HK68 immunized Bcl6^fl/fl^ or Bcl6^fl/fl^CD4^Cre^ mice to WT immunization-naïve mice (n=6 for each group) and a challenge with the X-31influenza strain. (F) Kaplan-Meier graph showing the probability of survival and (G) percentage of weight loss after lethal X-31 infection of WT mice with passive transfer of serum from Bcl6^fl/fl^, Bcl6^fl/fl^CD4^Cre^ mice or the transfer of PBS as a control. The humane endpoint post-viral challenge was 20% weight loss.

To confirm that protection was due to antibodies, the immune sera were collected from Bcl6^fl/fl^ and Bcl6^fl/fl^CD4^Cre^ mice (3 weeks after the 2^nd^ HK68 trimer immunization) and then transferred to unimmunized wild-type mice, which were then subjected to lethal X-31 virus challenge (Fig. 3E). The immune sera from the two genotypes were both protective [Bcl6^fl/fl^CD4^Cre^ immune sera (4 of 6) and Bcl6^fl/fl^ immune sera (5 of 6), each relative to non-immunized control (0 of 6), P<0.0024 and P<0.0061, respectively (Mantel-Cox test of survivorship)] (Fig. 3F-G), indicating that vaccine induced antibody protection against homologous viral challenge occurs both with and without GCs.

We next extended this paradigm to include vaccine protection against more diverse, heterologous influenza viral strains, as evaluated by H1ssF-elicited immunity against unmatched maA/Cal/09, a mouse-adapted form of the 2009 pandemic H1N1 (A/California/07/2009) (Fig. 4A) (*30–32*). H1ssF is a rationally designed ferritin nanoparticle display (8mer) of a trimeric H1 stem domain (A/New Caledonia/20/1999), a relatively conserved HA moiety, and generates antibody-dependent protection against heterosubtypic group 1 influenza A viruses following 3x sequential immunization in mice (*33–35*). The H1ssF immunization regimen elicited significant protection from heterologous H1N1 challenge in both genotypes [Bcl6^fl/fl^CD4^Cre^ mice (5 of 15) and Bcl6^fl/fl^ mice (8 of 15), each relative to non-immunized control (0 of 10), P<0.038 and P<0.0059, respectively (Mantel-Cox test of survivorship)] (Fig. 4B-D). Akin to our prior results with homologous H3N2 challenge, passive transfer of immune sera (3 weeks after the last H1ssF immunization) conferred significant protection from the same heterologous H1N1 challenge within naïve wildtype mice [Bcl6^fl/fl^CD4^Cre^ immune sera (4 of 10) and Bcl6^fl/fl^ immune sera (5 of 10), each relative to non-immunized control (0 of 10), P<0.0085 and P<0.0022, respectively (Mantel-Cox test of survivorship)] (Fig. 4E-G). Collectively, these data demonstrate that broad antibody-based protection against diverse lethal influenza viral challenges can also be elicited in the absence of GCs, potentiating a broadly protective activity of the non-GC memory B cell compartment.

**Figure 4.**
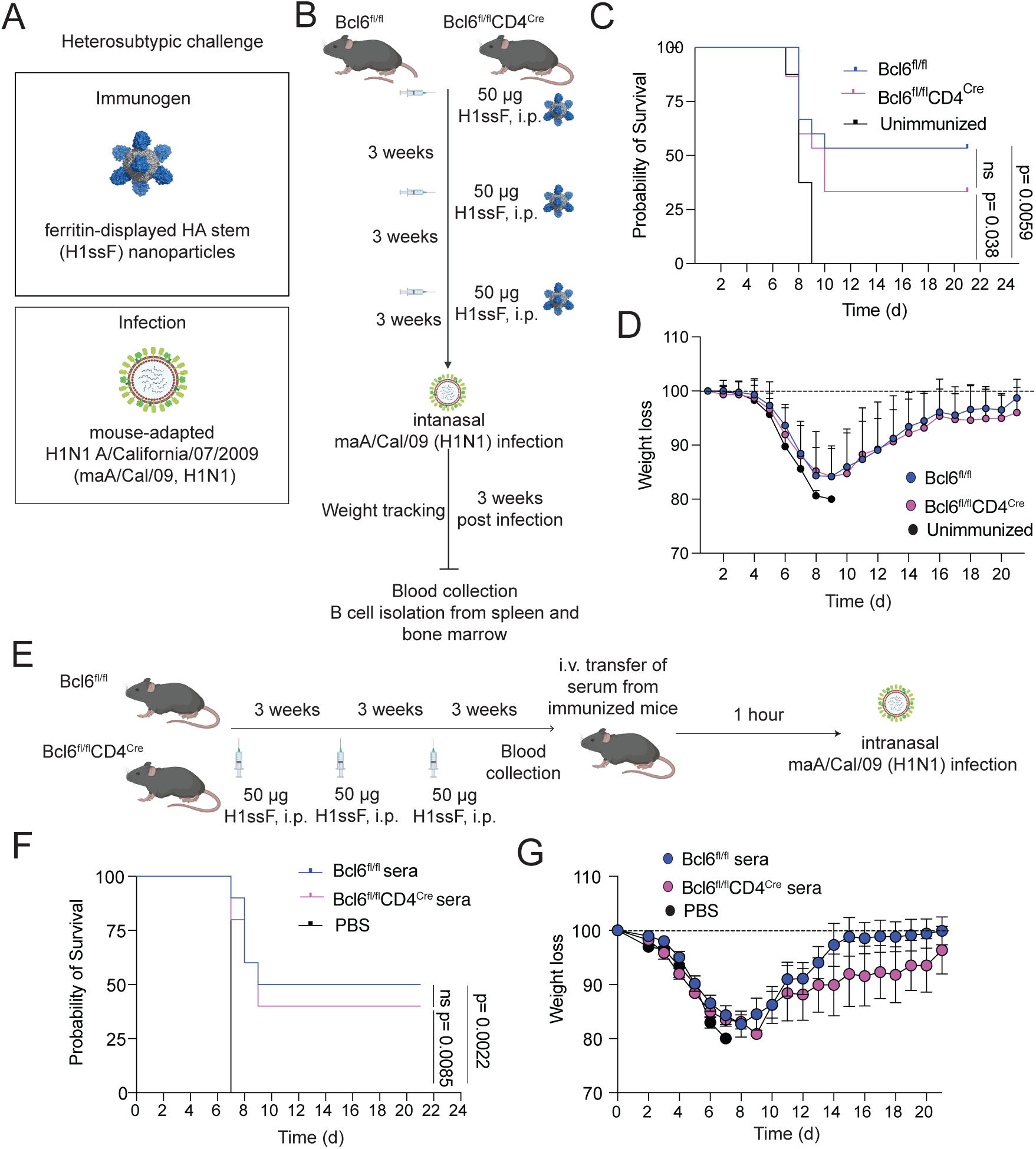
Non-GC derived humoral immunity protects against a lethal heterosubtypic influenza challenge. (A, B) 3 weeks after the 3^rd^ 50 µg dose of the H1ssF immunogen, Bcl6^fl/fl^ (n=15) and Bcl6^fl/fl^CD4^Cre^ (n=15) mice were challenged with a heterosubtypic influenza virus, maA/Cal/09 (H1N1). (C) A Kaplan-Meier graph showing the probability of survival and (D) the percentage of weight loss after maA/Cal/09 infection in immunized Bcl6^fl/fl^, Bcl6^fl/fl^CD4^Cre^ and unimmunized mice. Data are aggregated from two independent experiments. (E) Experimental plan for the passive transfer of sera from ssNP immunized Bcl6^fl/fl^ or Bcl6^fl/fl^CD4^Cre^ mice to WT immunization naïve mice (n=10 for each group) followed by a challenge with influenza maA/Cal/09. (F) Kaplan-Meier graph showing the probability of survival and (G) percentage of weight loss after mCal09 infection of WT mice following passive transfer of serum from Bcl6^fl/fl^ or Bcl6^fl/fl^CD4^Cre^ mice or the injection of PBS alone as a control. The humane endpoint post-viral challenge was 20% weight loss.

#### Vaccine-induced class-switched non-GC memory B cells directly protect from lethal inffuenza challenge

Our finding that HA-specific class-switched GC-independent memory B cells were significantly enriched in mice that survived the lethal challenge by the homologous and heterologous influenza viruses suggested a contribution of non-GC memory B cells to the antibody-dependent protection we observed (Fig. 5A-B, Supp, Fig. 6). To test this, we sorted class-switched GC-memory and non-GC memory B cells from wild-type mice immunized with HK68 trimers (collected at 3 weeks post-boost) and adoptively transferred 50-100K memory cells into µMt^-^ mice, which do not have a functional mature B cell compartment (Fig. 5C). Following lethal X-31 virus challenge, both GC-derived and non-GC memory B cells conferred protection in this model [non-GC-memory B cells (5 of 10) and GC-memory B cells (6 of 10), each relative to µMt control (0 of 10), P<0.0001 and P<0.0001, respectively (Mantel-Cox test of survivorship)] (Fig. 5C-E), demonstrating that the non-GC memory B cells enable rapid recall of influenza-protective extrafollicular antibodies.

**Figure 5.**
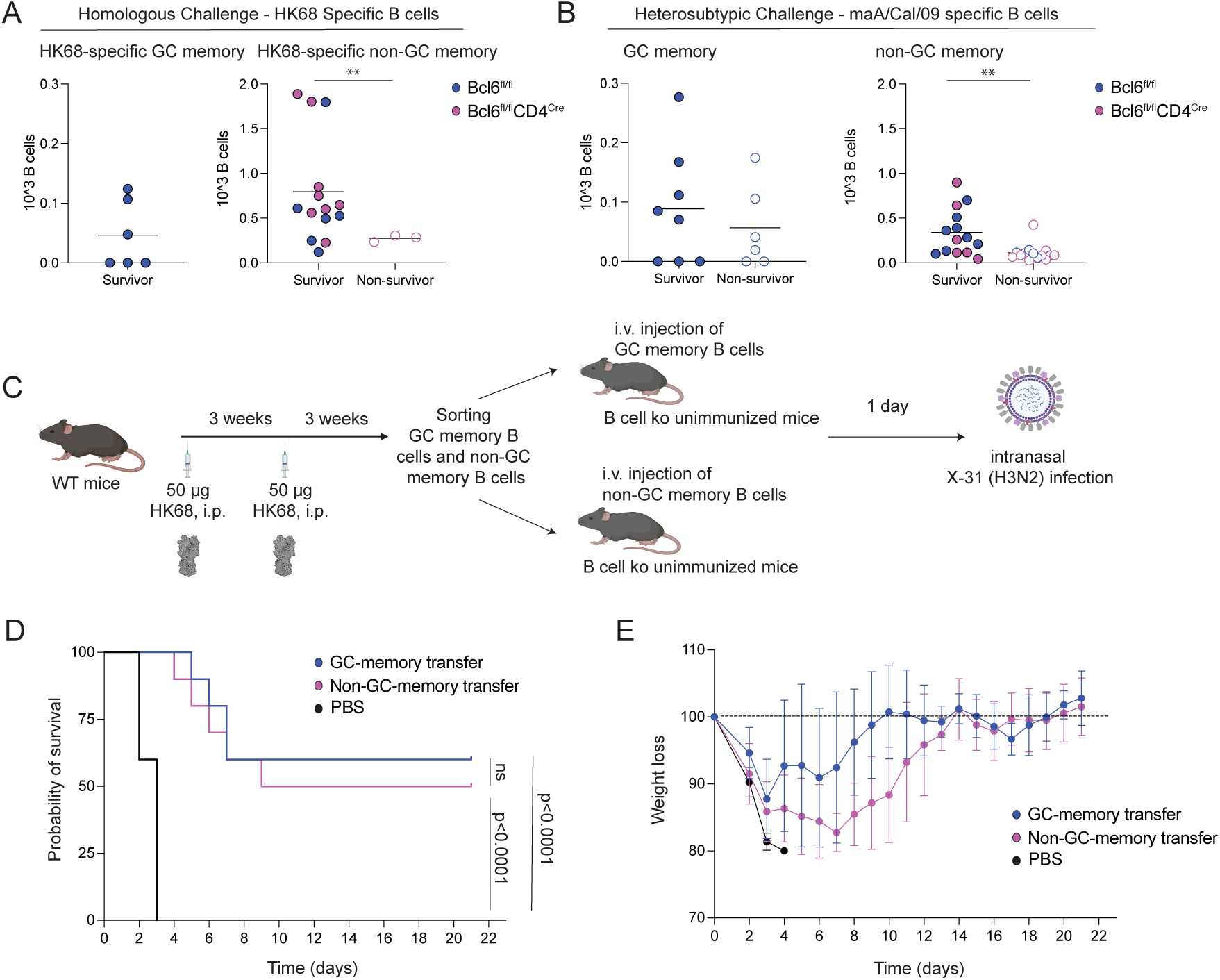
Vaccine-induced non-GC memory B cells provide rapid protection against lethal influenza challenge. (A) HK68-specific B cell numbers and (B) maA/Cal/09-specific B cell numbers for each non-GC resting B cell cluster in survivors, non-survivors and unimmunized (non-survivor) mice). Each dots represent a mouse. (C), GC-memory and non-GC-memory B cells were sorted 3 weeks after wild type mice were (n=60) immunized with two 50 µg doses of an HK68 derived hemagglutin trimer and these cells were separately transferred into μMt^-^ mice (lacking B cells) (n=10 for each group). 1 day after the transfer, recipient mice were challenged intranasally with a lethal dose of X-31 (H3N2) virus and monitored for up to 21 days post infection. (D) A Kaplan-Meier graph showing the probability of survival and (E) percentage of weight loss after X-31 challenge. The humane endpoint post-viral challenge was 20% weight loss.

## Discussion

Here we addressed two fundamental questions. Can durable class-switched memory B cell responses be generated in the absence of GCs? Can non-GC-derived class-switched memory B cells directly protect against lethal viral challenges?

It has long been established that the GC reaction leads to the generation of durable humoral immune responses and protective immunity in the context of vaccination and most infections (*1–3*). Severe infections, however, can result in the attenuation of GC formation and the skewing of humoral immunity towards extrafollicular B cell responses (*C*–*8*, *3C*, *37*). Although extrafollicular B cell populations have been widely studied in the context of severe infections, chronic inflammation and autoimmunity, class-switched non-GC-derived durable memory B cells have not been well characterized, and no studies using adoptive B cell transfer have formally established that class-switched memory B cells, whether GC-derived or generated in the course of an extrafollicular response, can offer protection from a lethal challenge (*14–17*, *38*).

In human studies, we and others have shown that resting extrafollicular memory (IgD^-^CD27^-^ CXCR5^+^) B cells have lower SHM levels and are as durable as CD27^+^ GC-derived switched memory B cells (*17*, *3S*–*41*). Nonetheless, it has not been possible to determine conclusively if these human putative non-GC switched memory B cells emerge outside GCs and, of course, in human studies, we lack the ability to definitively establish the functional relevance of specific lymphocyte populations unless there is a monogenic disease that eliminates them.

Our studies described here address this knowledge gap using an established genetically engineered mouse model that lacks Tfh cells and GCs. We have shown that two categories of durable class-switched memory B cells are generated in response to protein immunogens, including one population that can be generated in the absence of GCs. Non-GC memory B cells, defined as class switched Gl7^-^CD150^-^CD148^+^ B cells, have greater BCR diversity and lower SHM levels compared to GC-derived memory B cells and are largely generated by T-cell dependent responses. Our results indicate that class-switched non-GC memory B cells continued to expand for many months after vaccination, and are at least as durable as GC-derived memory B cells. Our studies also establish that humoral immune responses to influenza HA that are generated in the absence of GCs can protect recipient mice from a lethal challenge with matched and more heterologous influenza strains, underscored by a GC-independent memory B cell compartment that is positioned to generate rapid broadly protective humoral immunity against these pathogens. This humoral protection likely incorporates both neutralizing and non-neutralizing immunogloblin modalities, as have been described for antibodies targeting epitopes on HA (*34*, *42*–*4C*). A particularly interesting finding is that even in wild-type mice, the only class-switched B cell populations that correlate with survival against a homologous and heterosubtypic viral challenge were non-GC memory B cell subsets. In addition to their capacity to rapidly turn into antibody-secreting cells, the broader repertoires of non-GC memory B cells may better provide protection from life-threatening pathogens that rapidly generate variant surface antigens (*47*, *48*).

In conclusion, the studies described here provide definitive evidence that the GC is not the only source of durable class-switched memory B cells, and that class-switched non-GC B cells can, in isolation, provide protection from lethal pathogens. We conclude that while affinity maturation in the GC generates high-affinity antibodies, the non-GC class-switched memory B cell response preserves diversity and has the potential to provide rapid protection, particularly in the context of lethal and rapidly evolving pathogens.

## Methodology

### Mice

Bcl6^fl/fl^ [B6.129S(FVB)-^Bcl6tm1.1Dent^/J], Cd4^Cre^ [B6.Cg-Tg(Cd4-cre)1Cwi/BfluJ], RAG2 ko (B6.Cg-*Rag2^tm1.1Cgn^*/J), Tcrb ko (B6.129P2-*Tcrb^tm1Mom^*/J) and muMt^-^ (B6.129S2-*Ighm^tm1Cgn^*/J) mice were purchased from the Jackson Laboratory (Bar Harbor, ME). Bcl6^fl/fl^ mice were crossed with CD4^Cre^ mice to generate Bcl6^fl/fl^CD4^Cre^ mice. Mice were bred in-house using breeding pairs. For serum passive transfer and viral challenge studies, WT C57Bl/6J mice were also purchased from The Jackson Laboratory (Bar Harbor, ME). Mice of both sexes between 6 and 10 weeks old were utilized as littermates and cage pairs for this study. Sample sizes are indicated in the figure legends and were selected based on our orior studies on immunized mice that were validated by statistical analyses. For serum and cell transfer experiments, mice were randomized to age and sex matched groups. The investigators were not blinded. The mouse maintenance and experiments were performed following the approved protocols by the Institutional Animal Care and Use Committee (IACUC) of Massachusetts General Hospital (MGH), an Association for Assessment and Accreditation of Laboratory Animal Care International (AAALAC)–accredited facility, under Animal Study Protocol 2005N000360 and 2014N000252. Mice were genotyped by Transnetyx. The light cycles in the animal room were set on a 12-hour light cycle [7AM-7PM (ON) 7PM-7AM (OFF)]. The temperature range for the room was 68 to 73 degrees Fahrenheit, and the humidity index was from 30% to 70%. The feed was replaced every two weeks with fresh pelleted ration (Prolab Isopro RMH 3000), concomitant with changing fresh bedding in the cage. The cages were also inspected daily, and additional pellets were added if food was low or empty.

### Mice immunizations and viral challenges

Bcl6^fl/fl^ and Bcl6^fl/fl^CD4^Cre^ mice and tcrb-/- ko mice were immunized with 2 doses of 20 µg gp140, i.p, 3 weeks apart. WT mice were immunized with 2 doses of 100 µg NP-KLH (Biosearch technologies inc, # NC1125442), i.p., 3 weeks apart for transfer experiments. Bcl6^fl/fl^ and Bcl6^fl/fl^CD4^Cre^ mice were immunized with either 2 doses of 50 µg H3 HK68 i.p. and intranasally challenged with a 100% lethal dose of either H3N2 X-31 virus (BEI Resources, NR-3483; 10⁸ TCID₅₀/ml; (*4S*)) or immunized with 3 doses of 50 µg H1ssF and intranasally challenged with 100% lethal dose of mouse-adapted H1N1 A/California/07/2009 (maA/Cal/09; 10^4^ TCID_50_/ml; (*4S*)). After intranasal challenges, animals were monitored daily for 21 days for survival and body weight changes. Mice reaching a weight loss of 20% were humanely euthanized. Both viruses were cultured in MDCK cells and quantified by MDCK cell TCID_50_. maA/Cal/09 was kindly provided by Sabra Klein and Andrew Pekosz, John Hopkins University.

### Passive transfer of immune sera and viral challenge

To assess serum antibody protection activity, sera from mice immunized with either group 2 HK68 trimers (2x sequentially) or H1ssF (3x sequentially), were pooled and administered intravenously (tail vein, 100 ul per mouse) to WT C57BL/6J mice. Two hours post-transfer, mice were intranasally challenged with a 100% lethal dose of either H3N2 X-31 virus (BEI Resources, NR-3483; 10⁸ TCID₅₀/ml) or the mouse-adapted H1N1 A/California/07/2009 strain (maA/Cal/09; 10⁴ TCID₅₀/ml). Animals were monitored daily for 21 days for survival and body weight changes. Mice reaching a weight loss of 20% were humanely euthanized.

### Cell Transfer Experiments

Gl7^-^IgD^-^IgM^-^CD150^+^CD205^+^CD148^-^ live B cells and Gl7^-^IgD^-^IgM^-^CD150^-^CD205^-^CD148^+^ live B cells were sorted as GC-dependent and non-GC resting B cell populations. CD44^+^CXCR5^hi^PD1^hi^ live CD4^+^ T cells were sorted as follicular helper T cells (Tfh). All samples were sorted using FACSAria Fusion Cell Sorter. 50-100K cells of T_fh_ and corresponding resting B cell population was transferred to Rag2ko mice by tail injection. 24 hours after, mice were injected with NP-KLH, ip. 10 days after, serum samples were collected, mice were sacrificed and B cells were phenotyped from spleen and bone marrow. For vaccine-induced protection experiment, GC-memory and non-GC-memory B cells were sorted separately as described above and 50-100K cells were transferred to uMt^-^ mice by tail injection. 24 hours after, mice were challenged with lethal dose of intranasal X-31 infection. Mice weights were measured up to 21 days post infection. Mice reaching a weight loss of 20% were humanely euthanized, in accordance with predefined endpoints.

### Sample preparation for scRNA-seq experiments

For class-switched B cell libraries, B cells isolated from the spleens of Bcl6^fl/fl^ and Bcl6^fl/fl^CD4^Cre^ mice were stained with CD19, IgD, IgM, Gl7, CD73 and Gl7-streptavidin antibodies. Next, the cells were stained with anti-mouse hashtag antibodies 1-5 (BioLegend). After hashtag stainings, samples were pulled together as Bcl6^fl/fl^ and Bcl6^fl/fl^CD4^Cre^ groups and stained with Totalseq-C mouse universal cocktail and Totalseq-C Streptavidin-C0971 (both from BioLegend) following the manufacturer’s protocol. Gl7^-^IgD^-^ IgM^-^ live B cells were sorted in 0.5% PBS-BSA buffer using FACSAria Fusion Cell Sorter.

Sorted samples were encapsulated using Chromium GEM-X (10X Genomics). Gene expression (GEX), surface protein expression (antibody-derived tags, ADT) and BCR (VDJ) libraries were generated using the Chromium GEM-X Single Cell 5’ v3 Dual Index kit with feature barcode technology (10X Genomics) following the manufacturer’s protocol. Libraries were pooled at a 4:1:1 GEX:ADT:VDJ ratio and sequenced via paired-end reads on a NextSeq 2000 instrument with 4 100-cycle P4 kit (Illumina) in total to get good sequencing depth per cell.

### Protein expression, purification, and flow probe generation

Recombinant HA trimer ectodomains [(A/Hong Kong/1/1968 (H3N2) (HK68) and H1 Cal09 from A/California/07/2009(H1N1) (Cal09)], HIV-1 gp140 env, H1ssF, and ferritin-only nanoparticles were produced using established protocols (*22*, *32–35*, *50*, *51*). The proteins were expressed in Expi293 following transfection using the ExpiFectamine™ 293 Transfection Kit (Thermo Fisher Scientific). Supernatants were harvested five days post-transfection, clarified by filtration (VacuCap 8/0.2 μm filters, Pall Corporation), and then buffer-exchanged into PBS by a tangential flow filtration system (Pall Corporation) using a 10kDa filter (Cytiva, OS010T12). HA trimers and gp140 env were purified by Ni Sepharose affinity chromatography (20 mM imidazole wash, 500 mM imidazole elution), whereas H1ssF was affinity purified using Erythrina cristagalli lectin–immobilized resin (EY Laboratories) and eluted with 0.2 M lactose after a PBS wash. All proteins were further resolved and separated by size-exclusion chromatography (SEC) (ÄKTA pure, Cytiva), using the Superdex 200 Increase 10/300 column (HA trimers) or the Superose 6 Increase10/300 column (HIV-1 gp140 env and H1ssF and ferritin nanoparticles). Fluorescent B-cell probes were generated by amine-reactive labeling of ferritin-based constructs (AF488, AF594 or AF647), followed by re-purification by SEC; or by BirA-catalyzed biotinylation of Avi-tagged constructs [both HA trimers and gp140 contained a C-terminal foldon-Avi-His tag (*22*, *32*, *34*, *35*, *50–52*)], followed by conjugation to streptavidin-BV711 or streptavidin-PE-Cy7 (*50*). HA trimer probes also contained the Y98F receptor binding site mutation to prevent additional HA-based lectin activity for cell surface sialyl oligosaccharide (*53*). 0.4 ug protein per assay used for each probe.

### Flow Cytometry

EasySep Mouse Pan-B Cell isolation kit (Stem Cell) was used for B cell enrichment from splenocyte and bone marrow cells. EasySep Mouse CD4^+^ T cell isolation kit (Stem Cell) was used for CD4 T cell enrichment from splenocytes.

BD Horizon Brilliant Stain Buffer Plus was used in 1/100 ratio to improve the staining outcome. All the antibodies with the clones and dilutions are listed in Supp. Table 1. Cells were incubated with the staining mix including fluorescent B cell probes in 0.5% PBS-BSA FACS buffer, 30 min at 4C. All samples were recorded using BD FACSymphony A5.

### Enzyme-linked immunosorbent assay

To quantify antigen-specific IgG titers, 96-well plates were coated overnight at 4 °C with 200 ng per well of NP-KLH overnight at 4C. Plates were blocked for 1 h at room temperature with blocking buffer (1% BSA in PBS containing 0.01% Tween-20). Serum samples were serially diluted in PBS and incubated in plates for 2 hours at room temperature (seven dilutions, starting dilution 1:20). Following washing, plates were incubated for 1 h at room temperature with a secondary antibody isotypes such as anti-mouse IgG-horseradish peroxidase (HRP) (#GENA931V) from Sigma Millipore, anti-mouse IgG1-HRP (#1073-05), anti-mouse IgG2b-HRP (#1093-05), anti-mouse IgG2c-HRP (#1077-05), or anti-mouse IgG3-HRP (#1103-05) from SouthernBiotech or anti-mouse IgM-HRP (#550588) from BD Biosciences at 1:5,000 in PBS for 1 h at RT. Plates were developed with TMB Stabilized Chromogen (#34029) from Thermo Fisher Scientific, stopped with 3 N hydrochloric acid. Absorbances at 450 nm and 570 nm were determined with a plate reader (Synergy Neo2, BioTek). ELISA curves were calculated and analyzed using GraphPad Prism 10.5.0 (GraphPad). Pooled sera from mice 3 weeks after the last dose of immunizations were used as reference standards to calculate arbitrary units.

### Single-cell RNA, CITE-seq, and BCR-seq Analyses

Single-cell RNA-seq libraries were sequenced on an Illumina NextSeq 2000 (26 bp Read 1, 10 bp Index, 92 bp Read 2). Raw sequencing data for gene expression, V(D)J, and hashtag oligos (HTO) were processed using Cell Ranger (v9.0.1). Downstream analyses were performed in R (v4.4.2) using Seurat (v5.1.0). Gene expression, V(D)J, and HTO matrices were loaded as distinct assays in separate seurat objects for Spleen and Bone Marrow CD138^+^TACI^+^ Cells, only Bone Marrow CD138^+^TACI^+^ Cells, and Class-switched B cells.

Cells were filtered based on standard quality control metrics (>500 UMIs per cell, >250 unique genes, >0.8 log10GenesPerUMI, and <20% mitochondrial reads). HTO data was used to retain singlets only, resulting in 12,423 cells (Class-switched B cells) for downstream analysis. Gene expression matrices were normalized and highly variable features identified, excluding mitochondrial, ribosomal, and Ig genes during scaling and principal component analysis. The top principal components were used for UMAP dimensionality reduction, and graph-based clustering was applied to identify transcriptionally distinct B cell populations. The CITE based UMAP was generated using ADT features excluding hashtag ones. Dimensionality reduction and clustering were performed using the same workflow described above.

BCR clonotypes were assigned by mapping V(D)J assemblies to cell barcodes using the scRepertoire (v2.6.0) pipeline via the createHTOContigList method. Clonal expansion was visualized by overlaying clone size on UMAP embeddings, and clonal connectivity between cell subsets was assessed by tracking shared BCR sequences across clusters. For pseudobulk analysis, single-cell populations defined by UMAP clusters were aggregated using the AggregateExpression function in Seurat to generate sample-level count matrices. Differential expression testing on these pseudobulk matrices was performed with DESeq2 (v1.42.1), applying standard normalization and multiple testing correction procedures.

## Statistical tests

Statistical analyses were performed using GraphPad Prism version 10.4 and R. Normality was estimated using Shapiro-Wilk tests. Normally distributed data were compared using unpaired t-test and Welch’s correction when both populations don’t have similar SD. All statistical tests were two-tailed. Kaplan-Meier analysis was used for the probability of survival analysis after infections. Figures were prepared using Adobe Illustrator version 29.8.2, Flowjo GraphPad Prism, R and BioRender.com.

## Data availability

Full single-cell sequencing data generated in this study have been deposited in the NCBI Gene Expression Omnibus (GEO) prior to publication (GSE337878). The reference dataset used for sequence alignment was the refdata-gex-GRCm39-2024-A mouse transcriptome dataset, provided by 10× Genomics (10×genomics.com).

## Acknowledgements

The authors gratefully acknowledge the support of the Ragon Institute Vivarium and Ragon Institute Immunology Core-flow Cytometry Facility. The authors are grateful to D. Collins and J. Urbach for helpful discussion and advice. SP was supported by an HMS-Abbvie award, by by a grant from the Leducq foundation, by NIH U19 AI110495, UM1 AI1144295 and also a Ragon Institute Strategic Initiative to SP and DL. DL was also supported by NIH grants (R01AI155447, R01AI195539 P30AI060354, and R21AI193280) and BDW was supported by CHAVD grant (NIH UM1 AI144462).

## Author Contributions

EC and SP conceptualized the study. EC designed and performed experiments and data analysis with input from SP, BDW and DL. NI, ECL, EB and DM contributed to mice experiments and maintained mice colonies. CA and JS produced all recombinant protein immunogens and B cell probes and influenza viruses for this study. FANM and EC performed influenza vaccinations, viral challenges and passive transfers. AH and CVG analyzed scRNAseq and bulk RNA-seq data. SP, BDW and DL supervised the work and obtained the funding. EC, BDW and SP wrote the initial draft. All authors contributed to the final draft.

## References

1. J. G. Cyster, C. D. C. Allen, B Cell Responses: Cell Interaction Dynamics and Decisions. Cell 177, 524–540 (2019).

2. G. D. Victora, L. Mesin, Clonal and cellular dynamics in germinal centers. Curr. Opin. Immunol. 28, 90–96 (2014).

3. G. D. Victora, M. C. Nussenzweig, Germinal Centers. Annu. Rev. Immunol. 40, 413–442 (2022).

4. J. A. Roco, L. Mesin, S. C. Binder, C. Nefzger, P. Gonzalez-Figueroa, P. F. Canete, J. Ellyard, Q. Shen, P. A. Robert, J. Cappello, H. Vohra, Y. Zhang, C. R. Nowosad, A. Schiepers, L. M. Corcoran, K.-M. Toellner, J. M. Polo, M. Meyer-Hermann, G. D. Victora, C. G. Vinuesa, Class-Switch Recombination Occurs Infrequently in Germinal Centers. Immunity 51, 337–350.e7 (2019).

5. C. Viant, T. Wirthmiller, M. A. ElTanbouly, S. T. Chen, E. E. Kara, M. Cipolla, V. Ramos, T. Y. Oliveira, L. Stamatatos, M. C. Nussenzweig, Germinal center–dependent and – independent memory B cells produced throughout the immune response. J. Exp. Med. 218, e20202489 (2021).

6. N. Kaneko, H.-H. Kuo, J. Boucau, J. R. Farmer, H. Allard-Chamard, V. S. Mahajan, A. Piechocka-Trocha, K. Lefteri, M. Osborn, J. Bals, Y. C. Bartsch, N. Bonheur, T. M. Caradonna, J. Chevalier, F. Chowdhury, T. J. Diefenbach, K. Einkauf, J. Fallon, J. Feldman, K. K. Finn, P. Garcia-Broncano, C. A. Hartana, B. M. Hauser, C. Jiang, P. Kaplonek, M. Karpell, E. C. Koscher, X. Lian, H. Liu, J. Liu, N. L. Ly, A. R. Michell, Y. Rassadkina, K. Seiger, L. Sessa, S. Shin, N. Singh, W. Sun, X. Sun, H. J. Ticheli, M. T. Waring, A. L. Zhu, G. Alter, J. Z. Li, D. Lingwood, A. G. Schmidt, M. Lichterfeld, B. D. Walker, X. G. Yu, R. F. Padera, S. Pillai, Loss of Bcl-6-Expressing T Follicular Helper Cells and Germinal Centers in COVID-19. Cell 183, 143–157.e13 (2020).

7. Y. Duan, M. Xia, L. Ren, Y. Zhang, Q. Ao, S. Xu, D. Kuang, Q. Liu, B. Yan, Y. Zhou, Q. Chu, L. Liu, X.-P. Yang, G. Wang, Deficiency of Tfh Cells and Germinal Center in Deceased COVID-19 Patients. *Curr*. Med. Sci. 40, 618–624 (2020).

8. R. A. Elsner, C. J. Hastey, K. J. Olsen, N. Baumgarth, Suppression of Long-Lived Humoral Immunity Following Borrelia burgdorferi Infection. PLOS Pathog. 11, e1004976 (2015).

9. M. C. Woodruff, R. P. Ramonell, D. C. Nguyen, K. S. Cashman, A. S. Saini, N. S. Haddad, A. M. Ley, S. Kyu, J. C. Howell, T. Ozturk, S. Lee, N. Suryadevara, J. B. Case, R. Bugrovsky, W. Chen, J. Estrada, A. Morrison-Porter, A. Derrico, F. A. Anam, M. Sharma, H. M. Wu, S. N. Le, S. A. Jenks, C. M. Tipton, B. Staitieh, J. L. Daiss, E. Ghosn, M. S. Diamond, R. H. Carnahan, J. E. Crowe, W. T. Hu, F. E.-H. Lee, I. Sanz, Extrafollicular B cell responses correlate with neutralizing antibodies and morbidity in COVID-19. Nat. Immunol. 21, 1506–1516 (2020).

10. S. C. Eisenbarth, F. Batista, J. Cyster, R. Elsner, G. Kelsoe, F. E. Lund, S. Pillai, I. Sanz, M. Shlomchik, K.-M. Toellner, C. Vinuesa, N. Baumgarth, A roadmap for defining “extrafollicular” B cell responses. Immunity 58, 2627–2645 (2025).

11. D. Callahan, S. Smita, S. Joachim, K. Hoehn, S. Kleinstein, F. Weisel, M. Chikina, M. Shlomchik, Memory B cell subsets have divergent developmental origins that are coupled to distinct imprinted epigenetic states. Nat. Immunol. 25, 562–575 (2024).

12. S. A. Jenks, K. S. Cashman, E. Zumaquero, U. M. Marigorta, A. V. Patel, X. Wang, D. Tomar, M. C. Woodruff, Z. Simon, R. Bugrovsky, E. L. Blalock, C. D. Scharer, C. M. Tipton, C. Wei, S. S. Lim, M. Petri, T. B. Niewold, J. H. Anolik, G. Gibson, F. E.-H. Lee, J. M. Boss, F. E. Lund, I. Sanz, Distinct Effector B Cells Induced by Unregulated Toll-like Receptor 7 Contribute to Pathogenic Responses in Systemic Lupus Erythematosus. Immunity 49, 725–739.e6 (2018).

13. R. A. Elsner, M. J. Shlomchik, Germinal Center and Extrafollicular B Cell Responses in Vaccination, Immunity, and Autoimmunity. Immunity 53, 1136–1150 (2020).

14. S. Moir, J. Ho, A. Malaspina, W. Wang, A. C. DiPoto, M. A. O’Shea, G. Roby, S. Kottilil, J. Arthos, M. A. Proschan, T.-W. Chun, A. S. Fauci, Evidence for HIV-associated B cell exhaustion in a dysfunctional memory B cell compartment in HIV-infected viremic individuals. J. Exp. Med. 205, 1797–1805 (2008).

15. T.-A. Al-Aubodah, L. Aoudjit, G. Pascale, M. A. Perinpanayagam, D. Langlais, M. Bitzan, S. M. Samuel, C. A. Piccirillo, T. Takano, The extrafollicular B cell response is a hallmark of childhood idiopathic nephrotic syndrome. Nat. Commun. 14, 7682 (2023).

16. D. Y. Zhu, D. P. Maurer, C. Castrillon, Y. Deng, F. A. N. Mohamed, M. Ma, D. Tang, J. Min-Debartolo, N. Higginson-Scott, J. Buhlmann, A. G. Schmidt, D. Lingwood, M. C. Carroll, CD21 primes extrafollicular differentiation of autoreactive B cells in a TLR7-driven lupus model. Sci. Immunol. 10, eads8226 (2025).

17. C. A. Perugino, H. Liu, J. Feldman, J. Marbourg, T. V. Guy, A. Hui, N. Ingram, J. Liebaert, N. Chaudhary, W. Tao, C. Jacob-Dolan, B. M. Hauser, Z. Mian, A. Nathan, Z. Zhao, C. Kaseke, R. Tano-Menka, M. A. Getz, F. Senjobe, C. Berrios, O. Ofoman, Z. Manickas-Hill, D. R. Wesemann, J. E. Lemieux, M. B. Goldberg, K. Nündel, A. Moormann, A. Marshak-Rothstein, R. C. Larocque, E. T. Ryan, J. A. Iafrate, D. Lingwood, G. Gaiha, R. Charles, A. B. Balazs, A. Pandit, V. Naranbhai, A. G. Schmidt, S. Pillai, Two distinct durable human class-switched memory B cell populations are induced by vaccination and infection. Cell Rep. 44, 115472 (2025).

18. J. S. Chen, R. D. Chow, E. Song, T. Mao, B. Israelow, K. Kamath, J. Bozekowski, W. A. Haynes, R. B. Filler, B. L. Menasche, J. Wei, M. M. Alfajaro, W. Song, L. Peng, L. Carter, J. S. Weinstein, U. Gowthaman, S. Chen, J. Craft, J. C. Shon, A. Iwasaki, C. B. Wilen, S. C. Eisenbarth, High-affinity, neutralizing antibodies to SARS-CoV-2 can be made without T follicular helper cells. Sci. Immunol. 7, eabl5652 (2022).

19. K. Miyauchi, A. Sugimoto-Ishige, Y. Harada, Y. Adachi, Y. Usami, T. Kaji, K. Inoue, H. Hasegawa, T. Watanabe, A. Hijikata, S. Fukuyama, T. Maemura, M. Okada-Hatakeyama, O. Ohara, Y. Kawaoka, Y. Takahashi, T. Takemori, M. Kubo, Protective neutralizing influenza antibody response in the absence of T follicular helper cells. Nat. Immunol. 17, 1447–1458 (2016).

20. K. Hollister, S. Kusam, H. Wu, N. Clegg, A. Mondal, D. V. Sawant, A. L. Dent, Insights into the Role of Bcl6 in Follicular Th Cells Using a New Conditional Mutant Mouse Model. J. Immunol. 191, 3705–3711 (2013).

21. S. Crotty, T Follicular Helper Cell Biology: A Decade of Discovery and Diseases. Immunity 50, 1132–1148 (2019).

22. J. M. Kovacs, E. Noeldeke, H. J. Ha, H. Peng, S. Rits-Volloch, S. C. Harrison, B. Chen, Stable, uncleaved HIV-1 envelope glycoprotein gp140 forms a tightly folded trimer with a native-like structure. Proc. Natl. Acad. Sci. U. S. A. 111, 18542–18547 (2014).

23. G. Laszlo, K. S. Hathcock, H. B. Dickler, R. J. Hodes, Characterization of a novel cell-surface molecule expressed on subpopulations of activated T and B cells. J. Immunol. 150, 5252–5262 (1993).

24. K. M. Skrzypczynska, J. W. Zhu, A. Weiss, Positive Regulation of Lyn Kinase by CD148 Is Required for B Cell Receptor Signaling in B1 but Not B2 B Cells. Immunity 45, 1232– 1244 (2016).

25. A. H. Ellebedy, K. J. L. Jackson, H. T. Kissick, H. I. Nakaya, C. W. Davis, K. M. Roskin, A. K. McElroy, C. M. Oshansky, R. Elbein, S. Thomas, G. M. Lyon, C. F. Spiropoulou, A. K. Mehta, P. G. Thomas, S. D. Boyd, R. Ahmed, Defining antigen-specific plasmablast and memory B cell subsets in human blood after viral infection or vaccination. Nat. Immunol. 17, 1226–1234 (2016).

26. J. S. Chen, A. L. Tierney, O. A. Ojo, R. D. Chow, E. Song, T. Mao, B. Israelow, J. S. Weinstein, A. Williams, C. B. Wilen, S. C. Eisenbarth, Th1 cells express Tfh effector molecules and associate with B cells at the T-B border following viral infection. Immunology [Preprint] (2025). 10.1101/2025.04.02.646697.

27. S. T. Kim, J.-Y. Choi, B. Lainez, V. P. Schulz, D. E. Karas, E. D. Baum, J. Setlur, P. G. Gallagher, J. Craft, Human Extrafollicular CD4+ Th Cells Help Memory B Cells Produce Igs. J. Immunol. 201, 1359–1372 (2018).

28. J. L. Johnson, R. L. Rosenthal, J. J. Knox, A. Myles, M. S. Naradikian, J. Madej, M. Kostiv, A. M. Rosenfeld, W. Meng, S. R. Christensen, S. E. Hensley, J. Yewdell, D. H. Canaday, J. Zhu, A. B. McDermott, Y. Dori, M. Itkin, E. J. Wherry, N. Pardi, D. Weissman, A. Naji, E. T. L. Prak, M. R. Betts, M. P. Cancro, The Transcription Factor T-bet Resolves Memory B Cell Subsets with Distinct Tissue Distributions and Antibody Specificities in Mice and Humans. Immunity 52, 842–855.e6 (2020).

29. S. K. Lee, R. J. Rigby, D. Zotos, L. M. Tsai, S. Kawamoto, J. L. Marshall, R. R. Ramiscal, T. D. Chan, D. Gatto, R. Brink, D. Yu, S. Fagarasan, D. M. Tarlinton, A. F. Cunningham, C. G. Vinuesa, B cell priming for extrafollicular antibody responses requires Bcl-6 expression by T cells. J. Exp. Med. 208, 1377–1388 (2011).

30. A. L. Fink, K. Engle, R. L. Ursin, W.-Y. Tang, S. L. Klein, Biological sex affects vaccine efficacy and protection against influenza in mice. Proc. Natl. Acad. Sci. U. S. A. 115, 12477–12482 (2018).

31. R. L. Ursin, S. Dhakal, H. Liu, S. Jayaraman, H.-S. Park, H. R. Powell, M. L. Sherer, K. E. Littlefield, A. L. Fink, Z. Ma, A. L. Mueller, A. P. Chen, K. Seddu, Y. A. Woldetsadik, P. J. Gearhart, H. B. Larman, R. W. Maul, A. Pekosz, S. L. Klein, Greater Breadth of Vaccine-Induced Immunity in Females than Males Is Mediated by Increased Antibody Diversity in Germinal Center B Cells. mBio 13, e0183922 (2022).

32. R. Ray, F. A. Nait Mohamed, D. P. Maurer, J. Huang, B. A. Alpay, L. Ronsard, Z. Xie, J. Han, M. Fernandez-Quintero, Q. A. Phan, R. L. Ursin, M. Vu, K. H. Kirsch, T. Prum, V. C. Rosado, T. Bracamonte-Moreno, V. Okonkwo, J. Bals, C. McCarthy, U. Nair, M. Kanekiyo, A. B. Ward, A. G. Schmidt, F. D. Batista, D. Lingwood, Eliciting a single amino acid change by vaccination generates antibody protection against group 1 and group 2 influenza A viruses. Immunity 57, 1141–1159.e11 (2024).

33. H. M. Yassine, J. C. Boyington, P. M. McTamney, C.-J. Wei, M. Kanekiyo, W.-P. Kong, J. R. Gallagher, L. Wang, Y. Zhang, M. G. Joyce, D. Lingwood, S. M. Moin, H. Andersen, Y. Okuno, S. S. Rao, A. K. Harris, P. D. Kwong, J. R. Mascola, G. J. Nabel, B. S. Graham, Hemagglutinin-stem nanoparticles generate heterosubtypic influenza protection. Nat. Med. 21, 1065–1070 (2015).

34. M. Sangesland, L. Ronsard, S. W. Kazer, J. Bals, S. Boyoglu-Barnum, A. S. Yousif, R. Barnes, J. Feldman, M. Quirindongo-Crespo, P. M. McTamney, D. Rohrer, N. Lonberg, B. Chackerian, B. S. Graham, M. Kanekiyo, A. K. Shalek, D. Lingwood, Germline-Encoded Affinity for Cognate Antigen Enables Vaccine Amplification of a Human Broadly Neutralizing Response against Influenza Virus. Immunity 51, 735–749.e8 (2019).

35. M. Sangesland, A. Torrents de la Peña, S. Boyoglu-Barnum, L. Ronsard, F. A. N. Mohamed, T. B. Moreno, R. M. Barnes, D. Rohrer, N. Lonberg, M. Ghebremichael, M. Kanekiyo, A. Ward, D. Lingwood, Allelic polymorphism controls autoreactivity and vaccine elicitation of human broadly neutralizing antibodies against influenza virus. Immunity 55, 1693–1709.e8 (2022).

36. V. Ryg-Cornejo, L. J. Ioannidis, A. Ly, C. Y. Chiu, J. Tellier, D. L. Hill, S. P. Preston, M. Pellegrini, D. Yu, S. L. Nutt, A. Kallies, D. S. Hansen, Severe Malaria Infections Impair Germinal Center Responses by Inhibiting T Follicular Helper Cell Differentiation. Cell Rep. 14, 68–81 (2016).

37. R. A. Elsner, M. J. Shlomchik, IL-12 Blocks Tfh Cell Differentiation during Salmonella Infection, thereby Contributing to Germinal Center Suppression. Cell Rep. 29, 2796–2809.e5 (2019).

38. M. Gaya, K. L. Good-Jacobson, Molecular and tissue regulation of memory B cells. Sci. Immunol. 11, eaef3415 (2026).

39. Y. Zurbuchen, J. Michler, P. Taeschler, S. Adamo, C. Cervia, M. E. Raeber, I. E. Acar, J. Nilsson, K. Warnatz, M. B. Soyka, A. E. Moor, O. Boyman, Human memory B cells show plasticity and adopt multiple fates upon recall response to SARS-CoV-2. Nat. Immunol. 24, 955–965 (2023).

40. N. J. Bernard, Double-negative B cells. Nat. Rev. Rheumatol. 14, 684–684 (2018).

41. S. F. Andrews, M. J. Chambers, C. A. Schramm, J. Plyler, J. E. Raab, M. Kanekiyo, R. A. Gillespie, A. Ransier, S. Darko, J. Hu, X. Chen, H. M. Yassine, J. C. Boyington, M. C. Crank, G. L. Chen, E. Coates, J. R. Mascola, D. C. Douek, B. S. Graham, J. E. Ledgerwood, A. B. McDermott, Activation Dynamics and Immunoglobulin Evolution of Pre-existing and Newly Generated Human Memory B cell Responses to Influenza Hemagglutinin. Immunity 51, 398–410.e5 (2019).

42. D. J. DiLillo, G. S. Tan, P. Palese, J. V. Ravetch, Broadly neutralizing hemagglutinin stalk-specific antibodies require FcγR interactions for protection against influenza virus in vivo. Nat. Med. 20, 143–151 (2014).

43. D. J. DiLillo, P. Palese, P. C. Wilson, J. V. Ravetch, Broadly neutralizing anti-influenza antibodies require Fc receptor engagement for in vivo protection. J. Clin. Invest. 126, 605–610 (2016).

44. M. Sangesland, D. Lingwood, Antibody Focusing to Conserved Sites of Vulnerability: The Immunological Pathways for “Universal” Influenza Vaccines. Vaccines **G**, 125 (2021).

45. A. Watanabe, K. R. McCarthy, M. Kuraoka, A. G. Schmidt, Y. Adachi, T. Onodera, K. Tonouchi, T. M. Caradonna, G. Bajic, S. Song, C. E. McGee, G. D. Sempowski, F. Feng, P. Urick, T. B. Kepler, Y. Takahashi, S. C. Harrison, G. Kelsoe, Antibodies to a Conserved Influenza Head Interface Epitope Protect by an IgG Subtype-Dependent Mechanism. Cell 177, 1124–1135.e16 (2019).

46. A. L. Beukenhorst, K. L. Rice, J. Frallicciardi, M. H. Koldijk, C. M. Boudreau, J. Crawford, L. A. H. M. Cornelissen, K. A. S. da Costa, B. A. de Jong, S. Fischinger, B. Julg, J. M. Klap, C. M. Koch, Z. Magyarics, F. A. N. Mohamed, V. Okonkwo, L. Adams, C. M. McCarthy, L. Ronsard, N. Temperton, H. Vietsch, K. Wichapong, B. Ziere, D. Lingwood, J. Goudsmit, Intranasal administration of a panreactive influenza antibody reveals Fc-independent mode of protection. Sci. Rep. 15, 10309 (2025).

47. M. S. Miller, T. J. Gardner, F. Krammer, L. C. Aguado, D. Tortorella, C. F. Basler, P. Palese, Neutralizing Antibodies Against Previously Encountered Influenza Virus Strains Increase over Time: A Longitudinal Analysis. Sci. Transl. Med. 5 (2013).

48. D. H. Morris, V. N. Petrova, F. W. Rossine, E. Parker, B. T. Grenfell, R. A. Neher, S. A. Levin, C. A. Russell, Asynchrony between virus diversity and antibody selection limits influenza virus evolution. eLife 9, e62105 (2020).

49. R. Ray, F. A. Nait Mohamed, D. P. Maurer, J. Huang, B. A. Alpay, L. Ronsard, Z. Xie, J. Han, M. Fernandez-Quintero, Q. A. Phan, R. L. Ursin, M. Vu, K. H. Kirsch, T. Prum, V. C. Rosado, T. Bracamonte-Moreno, V. Okonkwo, J. Bals, C. McCarthy, U. Nair, M. Kanekiyo, A. B. Ward, A. G. Schmidt, F. D. Batista, D. Lingwood, Eliciting a single amino acid change by vaccination generates antibody protection against group 1 and group 2 influenza A viruses. Immunity 57, 1141–1159.e11 (2024).

50. G. C. Weaver, R. F. Villar, M. Kanekiyo, G. J. Nabel, J. R. Mascola, D. Lingwood, In vitro reconstitution of B cell receptor-antigen interactions to evaluate potential vaccine candidates. Nat. Protoc. 11, 193–213 (2016).

51. V. C. Rosado, L. Adams, A. S. Yousif, M. Sangesland, L. Ronsard, V. Okonkwo, C. McCarthy, C. Alexander, D. Irvine, D. Lingwood, A modular protocol for virosome display of subunit vaccine antigens. STAR Protoc. 6, 103610 (2025).

52. J. R. R. Whittle, A. K. Wheatley, L. Wu, D. Lingwood, M. Kanekiyo, S. S. Ma, S. R. Narpala, H. M. Yassine, G. M. Frank, J. W. Yewdell, J. E. Ledgerwood, C.-J. Wei, A. B. McDermott, B. S. Graham, R. A. Koup, G. J. Nabel, Flow cytometry reveals that H5N1 vaccination elicits cross-reactive stem-directed antibodies from multiple Ig heavy-chain lineages. J. Virol. 88, 4047–4057 (2014).

53. J. R. R. Whittle, R. Zhang, S. Khurana, L. R. King, J. Manischewitz, H. Golding, P. R. Dormitzer, B. F. Haynes, E. B. Walter, M. A. Moody, T. B. Kepler, H.-X. Liao, S. C. Harrison, Broadly neutralizing human antibody that recognizes the receptor-binding pocket of influenza virus hemagglutinin. Proc. Natl. Acad. Sci. U. S. A. 108, 14216– 14221 (2011).

