## Supplemental Data 1 for "Germinal center-independent memory B cells provide rapid protection from lethal influenza challenges": C╠oakan et al_supplementray materials.pdf

Supplementary materials include
Supplementary Figures 1-6
Supplementary Table 1

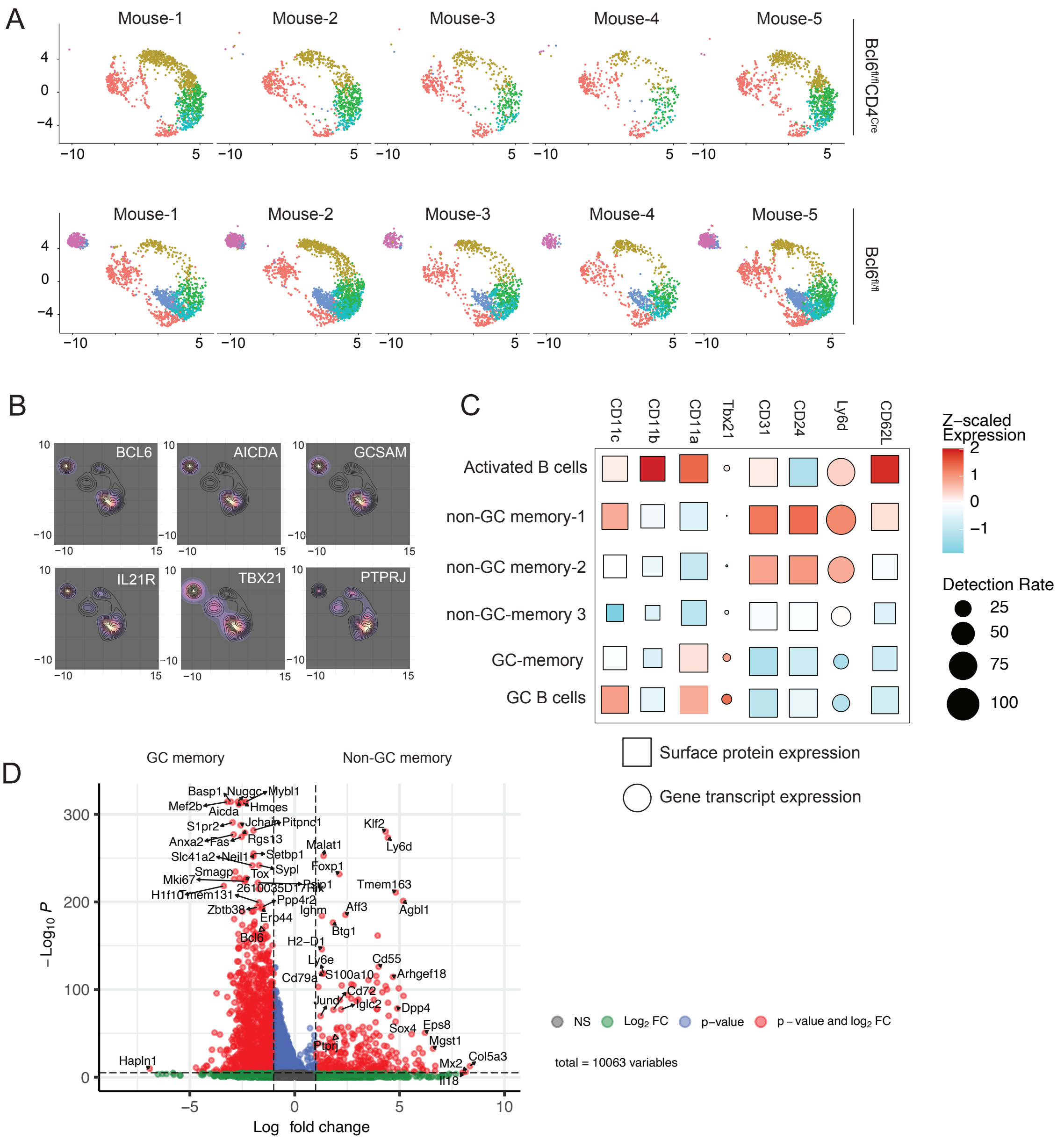


**Supp. Fig. 1.** GC linked B cells are not generated in Bcl6$^{fl/fl}$CD4$^{Cre}$ mice.

(A) UMAP-based clustering of IgD$^{-}$IgM$^{-}$ B cells for each mouse. (B) *Bcl6, Aicda, Gcsam, Tbx21, Zeb2, Ptprj* and *Il21r* gene expression in B cell clusters in Bcl6$^{fl/fl}$ mice (C) Bubble plots showing the average and percent expression of indicated genes in each cluster in Bcl6$^{fl/fl}$ (wild type) mice. (D) Volcano plot showing increased and decreased gene expression in non-GC memory B cells compared to GC-memory B cells.

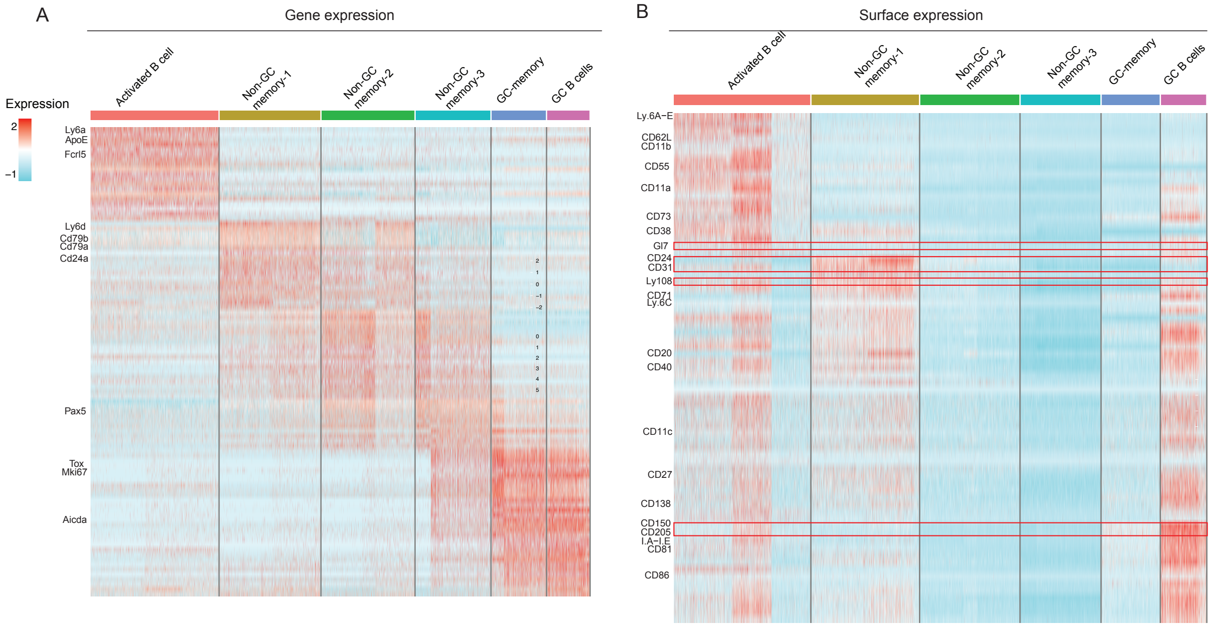


**Supp. Fig. 2**. Clustering of B cells in Bcl6$^{fl/fl}$ and Bcl6$^{fl/fl}$CD4$^{Cre}$ mice reveals GC dependent and independent B cell populations with different expression of surface markers and transcriptomic differences.

(A) Heat map generated using cluster-specific marker genes for IgD$^{-}$IgM$^{-}$ B cells isolated from Bcl6$^{fl/fl}$ (n=5) and Bcl6$^{fl/fl}$CD4$^{Cre}$ (n=5) mice 3 weeks post 2$^{nd}$ gp140 immunization and profiled using 5'-end droplet-based scRNA-seq. Columns in the heat map represent cells; rows represent cluster specific genes. (B) Heat map generated using cluster-specific surface markers for different populations as explained above. Columns in the heat map represent cells; rows represent cluster specific surface markers.

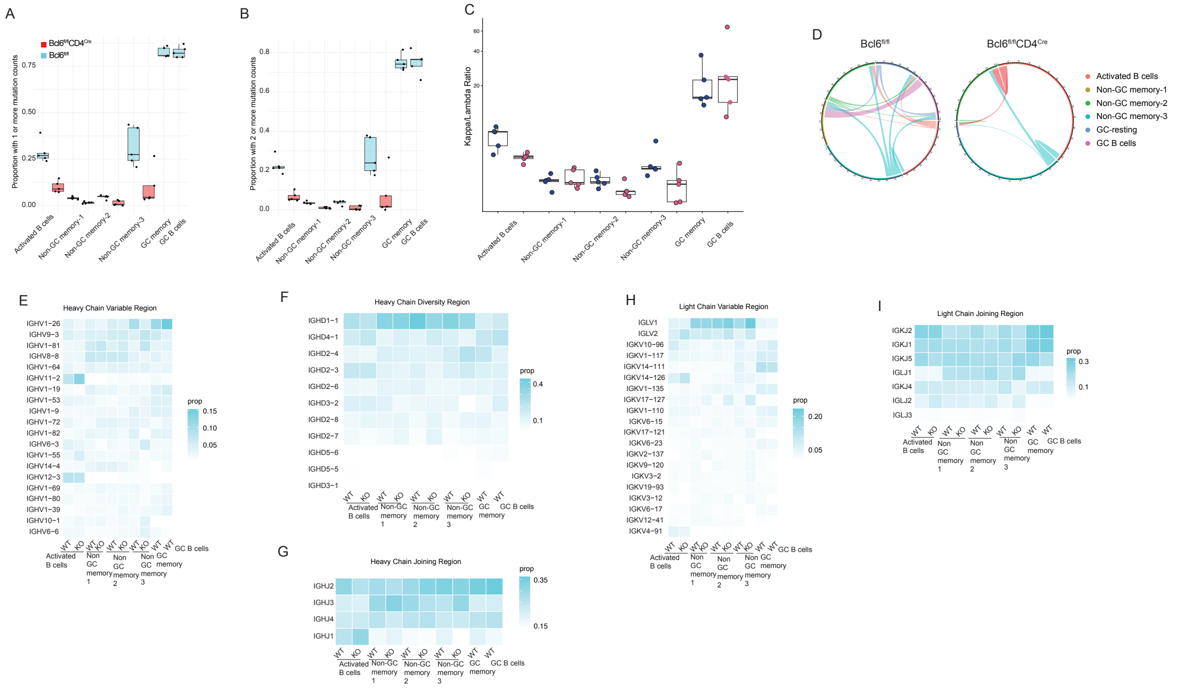


**Supp. Fig. 3.** Non-GC memory B cells exhibit low somatic hypermutation frequencies, different Ig gene segment usage during V(D)J recombination and have more diverse BCRs compared to GC-memory B cells.

(A) Proportion of cells with more than 1 mutation per rearranged Ig gene and (B) with more than 2 mutations are represented in each mouse for the indicated B cell clusters. Each dot represents a mouse. Box plots represent the mean, 1$^{st}$ and 3$^{rd}$ quartiles. (C) Kappa/lambda ratio for indicated clusters in Bcl6$^{fl/fl}$ or Bcl6$^{fl/fl}$CD4$^{Cre}$ mice. Each dot represents a mouse. (D) BCR connectivity between the clusters in Bcl6$^{fl/fl}$ and Bcl6$^{fl/fl}$CD4$^{Cre}$ mice. (F-I) Heatmaps showing the proportion of indicated V, D or J gene segments in heavy or light chains in Bcl6$^{fl/fl}$ (n=5) and Bcl6$^{fl/fl}$CD4$^{Cre}$ (n=5) mice.

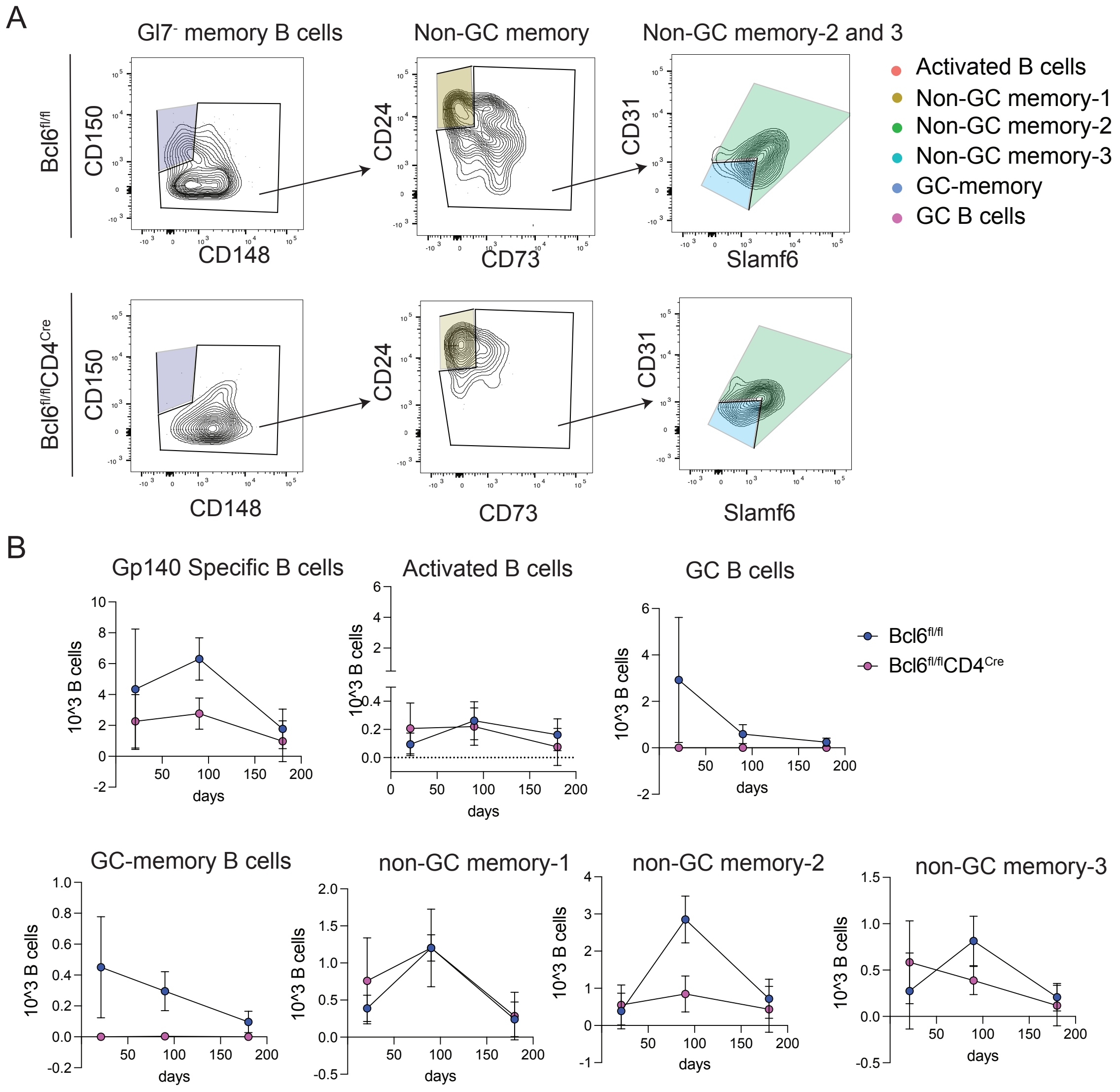


**Supp. Fig. 4.** Both GC and non-GC class switched B cells contribute to durable memory B cell populations.

(A) Gating strategy for dissecting gp140-specific memory B cells into class- switched GC-memory and 3 non-GC memory clusters using the cluster-defining markers generated from the surface expression library. (B) Total numbers of each gp140-specific B cell population 3 weeks, 3 months and 6 months after the 2$^{nd}$ immunization with gp140. Each dot represents the average of cell numbers in 5 mice.

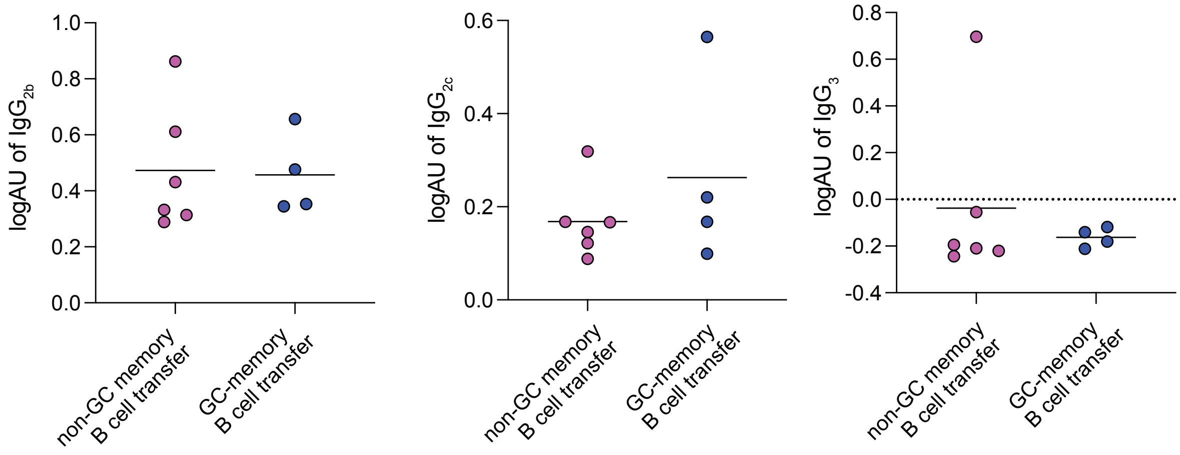


**Supp. Fig. 5**. Non-GC memory B cells contribute to memory recall responses and feed into new GC and non-GC reactions.

NP-KLH specific $IgG_{2b}$, $IgG_{2c}$ and $IgG_3$ serum levels at 10 days after NP-KLH immunization of Rag2 ko mice transferred with non-GC memory B cells and T follicular helper (Tfh) cells (n=6) or GC-memory B cell and Tfh cells (n=4).

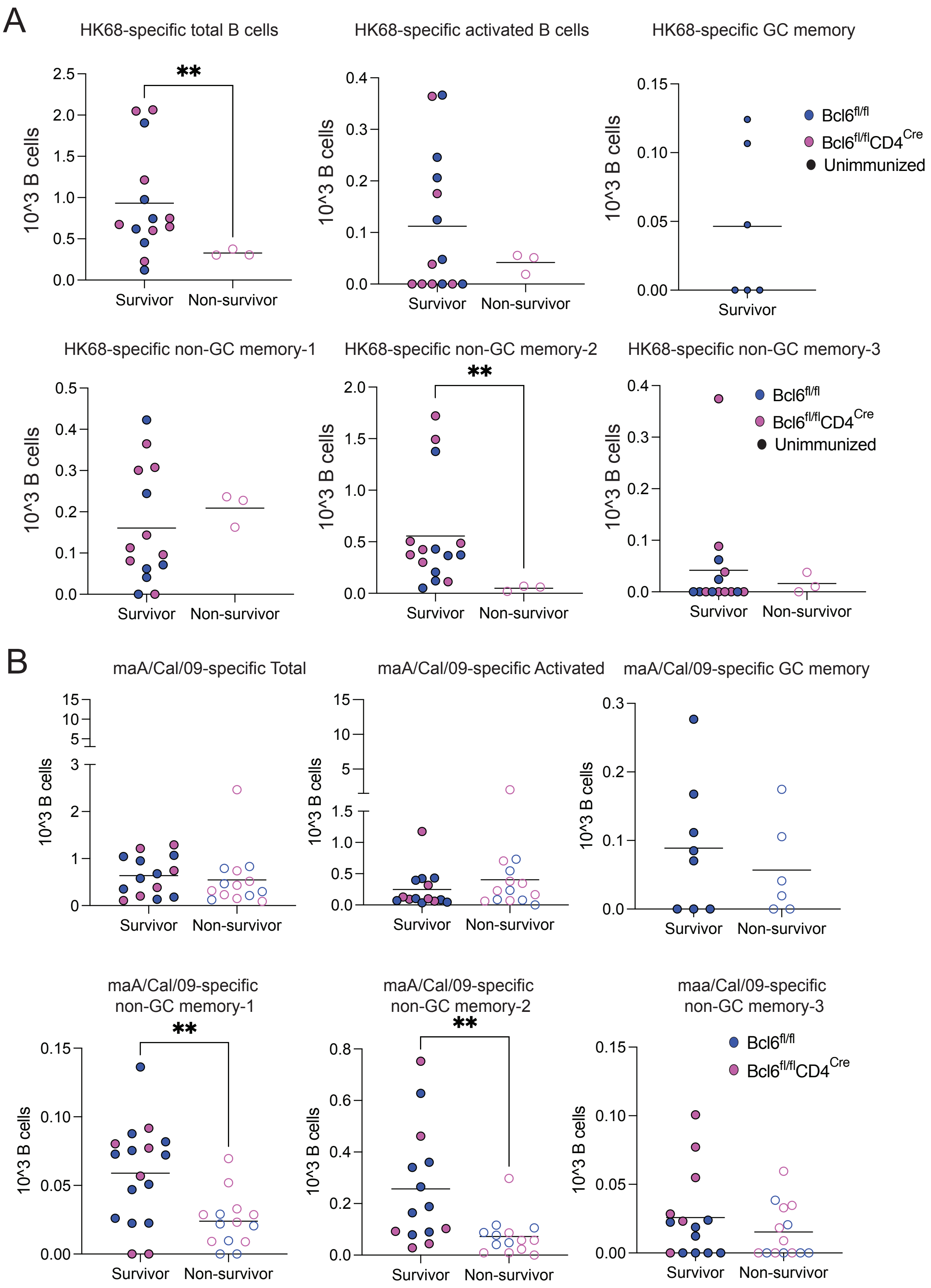
A
HK68-specific total B cells
HK68-specific activated B cells
HK68-specific GC memory
HK68-specific non-GC memory-1
HK68-specific non-GC memory-2
HK68-specific non-GC memory-3
10^3 B cells
Survivor
Non-survivor
Bcl6fl/fl
Bcl6fl/flCD4Cre
Unimmunized
B
maA/Cal/09-specific Total
maA/Cal/09-specific Activated
maA/Cal/09-specific GC memory
maA/Cal/09-specific non-GC memory-1
maA/Cal/09-specific non-GC memory-2
maa/Cal/09-specific non-GC memory-3

**Supp. Fig. 6.** Vaccine induced non-GC humoral immunity protects from homologous and heterologous influenza challenges.

(A) Numbers of HK-68 specific B cells from spleens of $Bcl6^{fl/fl}$ and $Bcl6^{fl/fl}CD4^{Cre}$ mice that were vaccinated with 2 doses of 50 μg i.p. H3 HK68 and challenged with a lethal dose of H3N2 X31 virus. Each dot represents a mouse. Mice reaching a weight loss of 20% were humanely euthanized, as "non-survivors". (B) The B cell numbers of maA/Cal/09-specific total or indicated clusters from spleens of $Bcl6^{fl/fl}$ and $Bcl6^{fl/fl}CD4^{Cre}$ mice vaccinated intraperitoneally with three 50 μg doses of H1ssF and challenged with a lethal dose of H1N1 maA/Cal/09 virus. Each dot represents a mouse.

**Supplementary Table 1**. Antibody reagents, dye and dilutions used in this study

| Supplier | Catalog No | Antigen | Fluor. or Conj. | Working Dilution |
|---|---|---|---|---|
| BioLegend | 144609 | GI7 | PerCp-Cy5.5 | 1:25 |
| BioLegend | 115506 | CD19 | FITC | 1:100 |
| BD Biosciences | 563157 | CD19 | BV711 | 1:100 |
| BD Biosciences | 562291 | CD19 | Pe-CD594 | 1:100 |
| BD Biosciences | 564274 | IgD | BUV395 | 1:50 |
| BioLegend | 405714 | IgD | APC | 1:50 |
| BD Biosciences | 115506 | CD4 | BUV395 | 1:100 |
| BD Biosciences | 563089 | CD31 | BV510 | 1:100 |
| BioLegend | 565747 | CD148 | PE | 1:100 |
| BD Biosciences | 756457 | IgM | RB780 | 1:25 |
| BioLegend | 134609 | Ly108 | APC | 1:50 |
| BD Biosciences | 565176 | CD138 | APC-R700 | 1:100 |
| BioLegend | 115926 | CD150 | BV421 | 1:50 |
| BD Biosciences | 612832 | CD24 | BUV737 | 1:100 |
| BD Biosciences | 749335 | CD24 | BUV805 | 1:100 |
| BioLegend | 127215 | CD73 | BV605 | 1:25 |
| Biolegend | 145511 | CXCR5 | BV421 | 1:50 |
| BioLegend | 135231 | PD-1 | BV711 | 1:200 |
| BioLegend | 103047 | CD44 | BV605 | 1:200 |
| BD Biosciences | 557653 | CD95 | PE-Cy7 | 1:200 |
| BioLegend | 107219 | CD273 (PD-L2) | BV421 | 1:100 |
| BD Biosciences | 741956 | CD80 | BUV805 | 1:25 |
| Fisher Scientific | L34976 | Invitrogen LIVE/DEAD Fixable Near-IR Dead Cell Stain | Near-IR | 1:1000 |
| BD Biosciences | 553142 | CD16/CD32 (Fc block) | | 1:100 |
| BD Biosciences | 405207 | Streptavidin | APC | |
| BD Biosciences | 405203 | Streptavidin | PE | |
| BioLegend | 405206 | Streptavidin | PE-Cy7 | |
| BD Biosciences | 563262 | Streptavidin | BV711 | |
